# Metabolic changes to host cells with *Toxoplasma gondii* infection

**DOI:** 10.1101/2023.08.10.552811

**Authors:** Gina M. Gallego-López, Emmanuel Contreras Guzman, Laura J. Knoll, Melissa Skala

## Abstract

*Toxoplasma gondii*, the causative agent of toxoplasmosis, is an obligate intracellular parasite that infects warm-blooded vertebrates across the world. In humans, seropositivity rates of *T. gondii* range from 10% to 90%. Despite its prevalence, few studies address how *T. gondii* infection changes the metabolism of host cells. Here, we investigate how *T. gondii* manipulates the host cell metabolic environment by monitoring metabolic response over time using non-invasive autofluorescence lifetime imaging of single cells, seahorse metabolic flux analysis, reactive oxygen species (ROS) production, and metabolomics. Autofluorescence lifetime imaging indicates that infected host cells become more oxidized and have an increased proportion of bound NAD(P)H with infection. These findings are consistent with changes in mitochondrial and glycolytic function, decrease of intracellular glucose, fluctuations in lactate and ROS production in infected cells over time. We also examined changes associated with the pre-invasion “kiss and spit” process using autofluorescence lifetime imaging, which similarly showed a more oxidized host cell with an increased proportion of bound NAD(P)H over 48 hours. Glucose metabolic flux analysis indicated that these changes are driven by NADH and NADP+ in *T. gondii* infection. In sum, metabolic changes in host cells with *T. gondii* infection were similar during full infection, and kiss and spit. Autofluorescence lifetime imaging can non-invasively monitor metabolic changes in host cells over a microbial infection time-course.

## INTRODUCTION

Host cells have evolved elaborate systems to counteract pathogen invasion, establishment, and replication, including phagolysosomal fusion, reactive oxygen species (ROS), nitrogen intermediates, sequestration of nutrients, and apoptosis [1–3]. However, host cell metabolic response to a microorganism infection remains unclear. Here, we examine how *T. gondii* infection changes the host metabolism including redox balance and NAD(P)H binding activities using autofluorescence lifetime imaging of single cells over an infection time-course, along with metabolic flux analysis, ROS production, metabolite sensing, and glucose labeling.

Previous studies have investigated how *T. gondii* infection reprograms the host cell. Blader *et al*., classified host genes modulated in response to *T. gondii* infection into three functionality different classes: (i) genes required for host defense; (ii) genes require parasite growth; and (iii) genes incidentally regulated as a consequence of modulating the first two classes [4]. Quantitative proteomic studies have suggested a global reprogramming of the cell metabolism by the parasite [1]. We recently published how a *T. gondii* full infection [5] and the pre-invasion “kiss and spit” process [6] produces significant changes in host metabolites. How *T. gondii* infection changes host metabolism has not yet been examined by live single-cell imaging or seahorse to determine the metabolic changes that *T. gondii* infection induces over time.

Here, we use fluorescence lifetime imaging microscopy (FLIM) of the autofluorescent metabolites NADH, NADPH, and FAD non-invasively monitor single host cell response to *T. gondii* infection over 48 hours. NADH and NADPH have overlapping fluorescence properties and are collectively referred to as NAD(P)H [7,8]. FLIM of NAD(P)H and FAD, or optical metabolic imaging (OMI), measures the optical redox ratio (ORR) defined as the fluorescence intensity of NAD(P)H / (FAD + NAD(P)H) (Table 1) [9–12]. ORR is an indicator of the oxidation-reduction state of the cell, and an important marker of cell health that can be used to monitor living tissues and cells. The ORR has been used to study numerous biological processes including cancer, thermal stress, de novo fatty acid synthesis, and diabetes [9–11,13]. Many factors can change the ORR, such as hypoxia, high carbon demands, increased proliferation rate, and fatty acid synthesis [11]. ORR imaging has been previously used in infectious disease research to monitor oxidative stress in host cells with chronic infection with hepatitis C virus (HCV) [14].

**Table 1:** OMI parameters and definitions.

| Parameters | Description |  |
| --- | --- | --- |
|  | NAD(P)H | FAD |
| $\tau_1$ | Free lifetime | Protein-bound lifetime |
| $\tau_2$ | Protein-bound lifetime | Free lifetime |
| $\alpha_1$ | Proportion of free NAD(P)H | Proportion of bound FAD |
| $\alpha_2$ | Proportion of bound NAD(P)H | Proportion of free FAD |
| $I$ | Absolute Intensity | Absolute Intensity |
| $\tau_m$ | Mean lifetime = $\alpha_1 \tau_1 + \alpha_2 \tau_2$ | |
| Optical Redox Ratio | $ORR = I_{NAD(P)H} / (I_{FAD} + I_{NAD(P)H})$ | |

OMI also provides a measurement of protein-binding activity for NAD(P)H and FAD [15,16]. Specifically, NAD(P)H has a short lifetime in the free conformation (τ_1_) and a long lifetime in the protein-bound conformation (τ_2_), while the converse is true for FAD (τ_1_ is bound, τ_2_ is free). Due to these distinct lifetimes, OMI can quantify the relative fractions of free and protein-bound NAD(P)H and FAD in the cell (Table 1) [17]. These fluorescence lifetimes have been previously used in numerous studies including infectious disease studies of HCV [18], *Chlamydia trachomatis* infection [19], and *Plasmodium falciparum* replication [20].

This study uses OMI to investigate how intracellular *T. gondii* manipulate their host cell metabolism. Human foreskin fibroblasts (HFF) cells were imaged with OMI over 48 hours of infection with *T. gondii.* Additional measurements of metabolic flux analysis, ROS production, intracellular and extracellular glucose and lactate provide validation of live cell OMI results. Comparisons of *T. gondii* full infection and the pre-invasion “kiss and spit” process were similarly performed with OMI. We aim to answer the following research questions: Does *T. gondii* infection affect the redox balance of the host cell? Does *T. gondii* infection affect the NAD(P)H or FAD binding activities of the host cell? Does OMI assess *T. gondii* infection consistently with standard measurements of cell metabolism? This research highlights the key role of unexplored redox biology in *T. gondii* infection.

## RESULTS

### Image analysis to quantify intracellular T. gondii

We established *T. gondii* infection in quiescent HFF cells using ME49 mCherry-labeled parasites because mCherry fluorescence is spectrally separate from both NAD(P)H and FAD fluorescence [21]. FAD and NAD(P)H intensities and lifetimes of infected cells were obtained using two-photon FLIM. OMI was collected at 1, 6, 9, 12, 24 and 48-hours post-infection (HPI) in two independent experiments. We used classical image processing techniques to create the host cell masks and the *T. gondii* masks (Figure 1). We used NAD(P)H images processed in CellProfiler and Napari to obtain the host cell masks, and *T. gondii* masks were created by processing the mCherry fluorescence images in python (Figure 1). After creating the final host cell masks and the final *T. gondii* masks for all the datasets, both masks were loaded into python to quantify the amount of *T. gondii* in each cell according to their overlap as shown in Figure 1.

**Figure 1.**
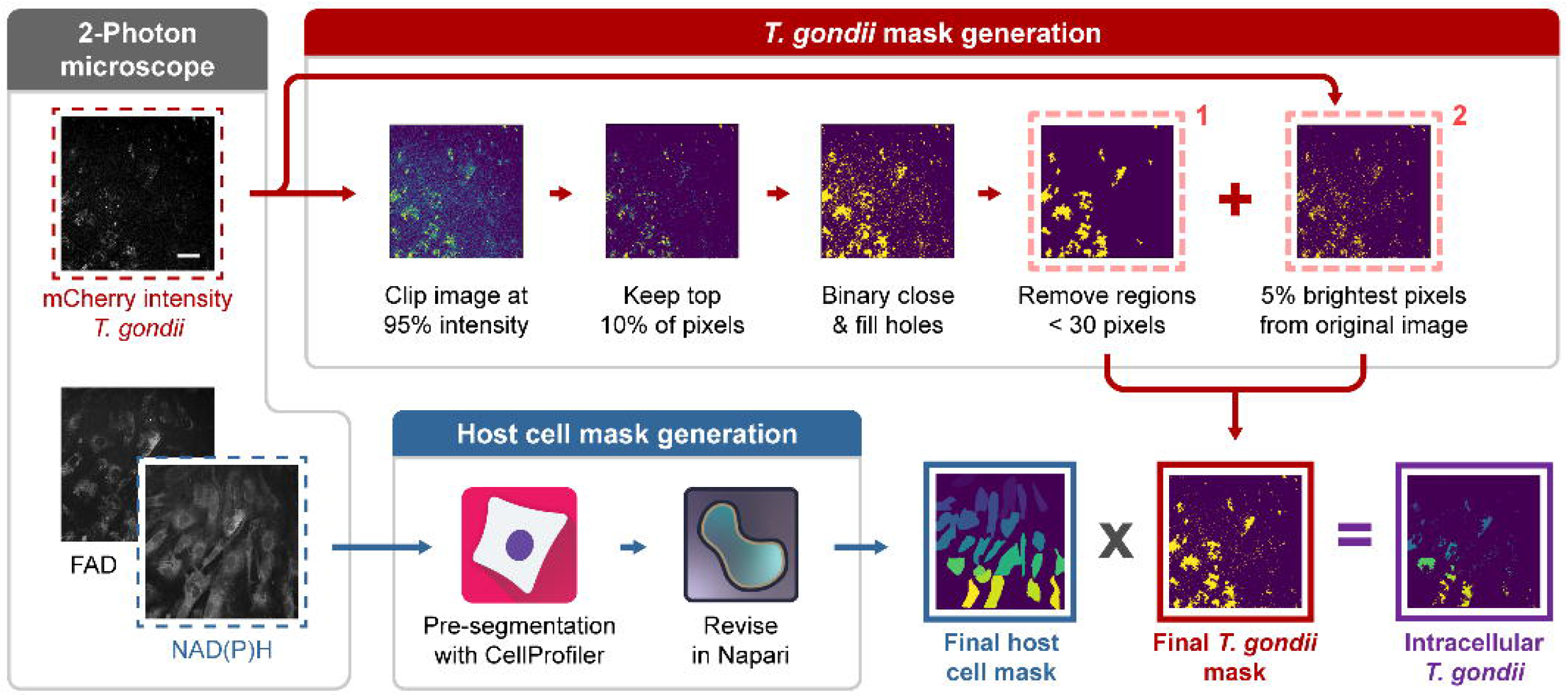
Pipeline for creating HFF host whole cell and *T. gondii* masks from two-photon FLIM images. For each field of view three images were taken: NAD(P)H (750 ex, 440/80 em.) and FAD (890 ex, 550/100 em.) intensity and lifetime as well as mCherry *T. gondii* (1090 ex, 690/50 em.) intensity. Scale bar = 50 µm. NAD(P)H intensity images were used to generate individual host cell masks with semi-automated segmentation methods using CellProfiler and manual revision in Napari. *T. gondii* masks were generated from the mCherry intensity images in Python using the package scikit-image (see methods).After creating the host whole cell masks and the *T. gondii* masks, both masks were used to quantify the amount of *T. gondii* in each cell according to their pixel overlap. This was done by multiplying the *T. gondii* mask by the host cell mask to produce a mask capturing the intersection of the parasite in each cell area.

### Establishing a T. gondii infection threshold

The percentage of intracellular *T. gondii* area by cell area was defined (Figure 2A). Variations in percent intracellular parasite showed that *T. gondii* did not infect quiescent HFF cells equally. To compare the infection distribution across the cells, we plotted histograms of percent intracellular *T. gondii* in each individual cell for each timepoint, for each experiment (Figure 2B, S1, S2, S3). The histogram in Figure 2B summarizes the percentage of intracellular *T. gondii* across two independent time course experiments. Figure S1 shows the percentage of intracellular *T. gondii* per timepoint in each independent experiment. Figure S2 and Figure S3 indicate the percentage of intracellular parasite in each time point in experiment #1 and #2, respectively.

**Figure 2.**
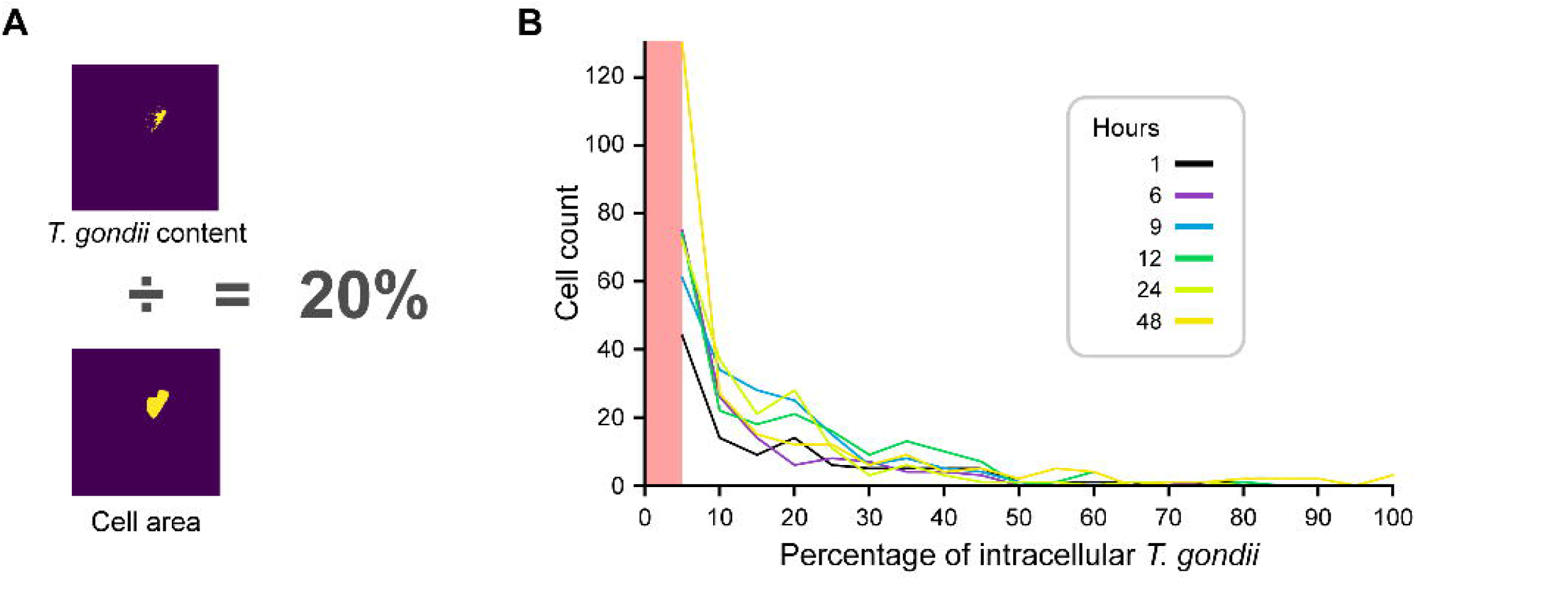
Percentage of intracellular *Toxoplasma gondii* infection per cell during time-course. **(A)** The percentage of intracellular *T. gondii* per cell was calculated by dividing the sum of the pixels in the mask belonging to *T. gondii* inside the cell by the number of pixels in the whole cell mask. **(B)** The percentage of intracellular parasite pixels per area of cell in 5% bins. The histogram shows the average of two independent experiments. X-axis shows the *T. gondii* percent area per cell. The Y-axis shows the cell count. The six different time points (in hours) are represented by different colors. The red box indicates cells with ≤5% *T. gondii* percent area per cell, which were excluded from the high *T. gondii* condition.

Given that not all quiescent HFF cells were equally infected by the parasite, we grouped cells into low and high infection categories initially by setting a threshold on the percent intracellular *T. gondii*. Due to relatively low signal-to-noise ratio of the mCherry labeled *T. gondii* images, some of the noise pixels in the masks have been erroneously quantified as parasite and thus we compared 5% and 10% infection thresholds to prevent cells from being quantified as false positive for containing the parasite. We then compared OMI parameters but did not see any significant differences in the results between the 5 and 10% thresholds (data not shown). We selected 5% as our cutoff threshold because of the similar OMI parameter results and because it divided our data into roughly half between the two infection categories. We used this analysis to empirically determine that cells with lower than 5% had no significant *T. gondii* infection or the infection could be a false positive due to pixel noise captured by the *T. gondii* masks.

### OMI changes in low vs high infected cells

Having categorized cells in the infected condition, we then compared low vs high parasite infected cells. OMI parameters were calculated as detailed in Figure 3. Figure 4A shows representative images outlining cells with low versus high infection with a 5% threshold. We used the fluorescence intensities of NAD(P)H and FAD to determine the ORR, which provides a label-free method to monitor the oxidation-reduction state of the cell (Figure 4B) [12]. Multiple definitions of the ORR exist, but here we used NAD(P)H/(NAD(P)H + FAD), (Table 1, and Figure 4B) as an increase in the ORR corresponds with a more reduced intracellular environment, suggestive of an increase in glycolysis, and it normalizes the values to be between 0 and 1 [22]. The decrease in ORR means a more oxidized intracellular environment, likely due to a decrease in glycolysis [23]. When comparing ORR, we observed a significant difference between low and high infection at 1 through 24 HPI (Figure 4C and Figure S4C). Cells with high infection showed a more oxidized ORR compared to cells with low infection. We did not find significant differences for NAD(P)H α_2_ or NAD(P)H τ_m_ between low and high categories besides NAD(P)H τ_m_ at the 9-, 24-, and 48-hour time points (Figure 4D, 4E, S4D and S4E).

**Figure 3.**
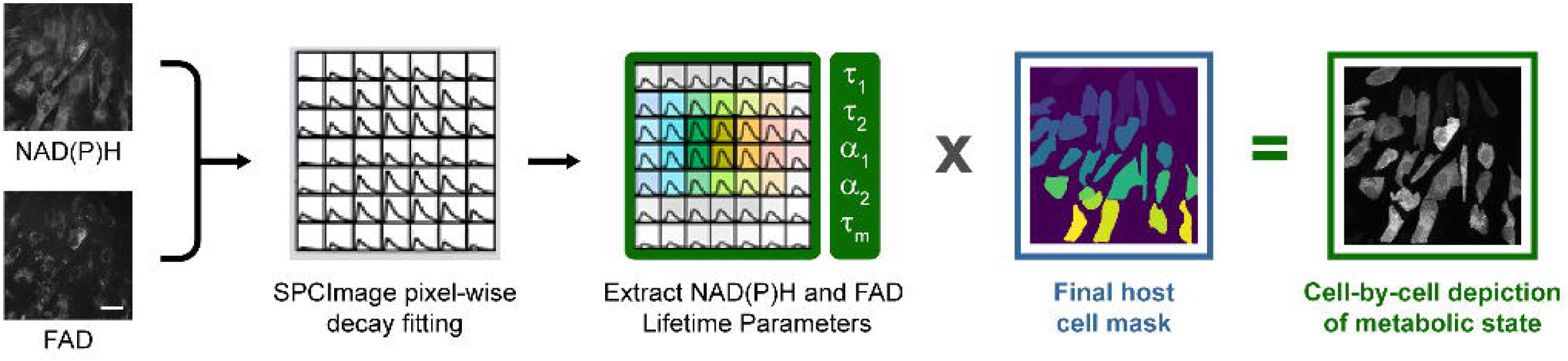
Standard analysis pipeline for calculating OMI parameters in whole cells. NAD(P)H and FAD lifetime images were used to calculate OMI parameters by fitting the decay curves in each pixel using SPCImage. Parameters were then averaged across each whole cell for NAD(P)H and FAD parameters using host cell masks generated in Figure 1, resulting in single cell depiction of metabolic state. Scale bar = 50 µm.

**Figure 4.**
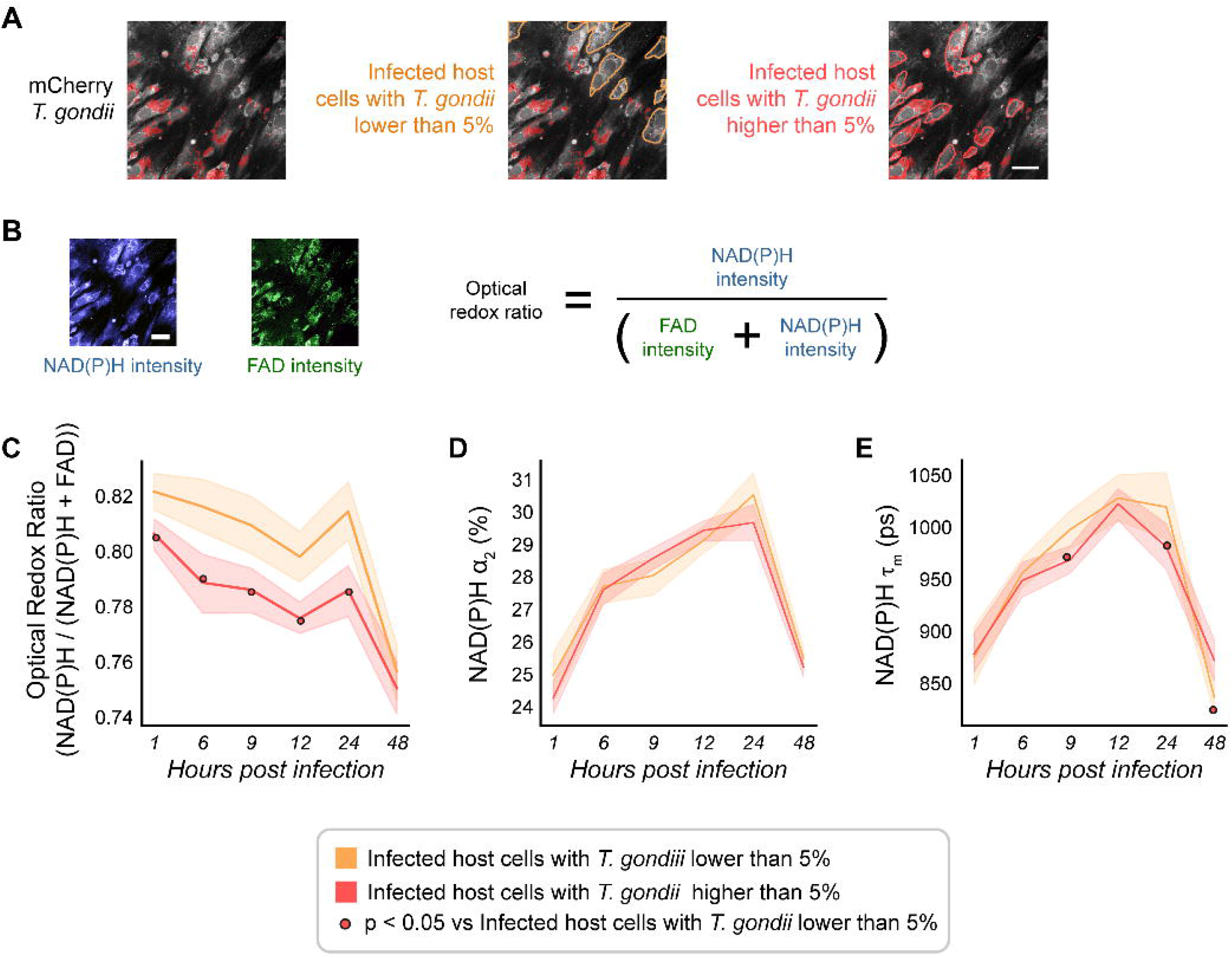
Establishing a threshold of *T. gondii* infection. **(A)** Representative images of infected HFF with lower than 5% (yellow) and higher (red) than 5% mCherry *T. gondii*. Scale bar = 50 µm. **(B)** Optical Redox ratio [fluorescence intensity of NAD(P)H / (FAD + NAD(P)H)]**. (C)** Optical Redox ratio of *T. gondii* infected HFF cells with low vs. high intracellular *Toxoplasma gondii* over time course of infection. **(D)** Percentage of protein bound NAD(P)H (α_2_) of *T. gondii* infected HFF cells with low vs. high intracellular *Toxoplasma gondii* over time course of infection. **(E)** Mean lifetime of NAD(P)H (τ_m_) of *T. gondii* infected HFF cells with low vs. high *Toxoplasma gondii* over time course of infection. The percentage of intracellular mCherry *T. gondii* lower than 5% is represented in yellow and higher than 5% is represented in red. Curves represent the mean and 95% confidence interval. Statistical significance was determined by an independent Student’s T-test. *p < 0.05. Cell count low *T. gondii* = 632, cell count high *T. gondii* = 974.

### T. gondii infection changes the redox balance of the host cell

Using our previously established intracellular parasites threshold of 5% to differentiate cells with low versus high infection, we compared uninfected HFF cells with high ME49 *T. gondii* infected cells. *T. gondii* infected cells exhibited a significant decrease in ORR compared to uninfected cells, suggesting a more oxidized microenvironment (Figure 5A and Figure S5A). Single cell analysis allows us to quantify these redox changes (Figure 5B). We found that infected cells become more oxidized over time.

**Figure 5.**
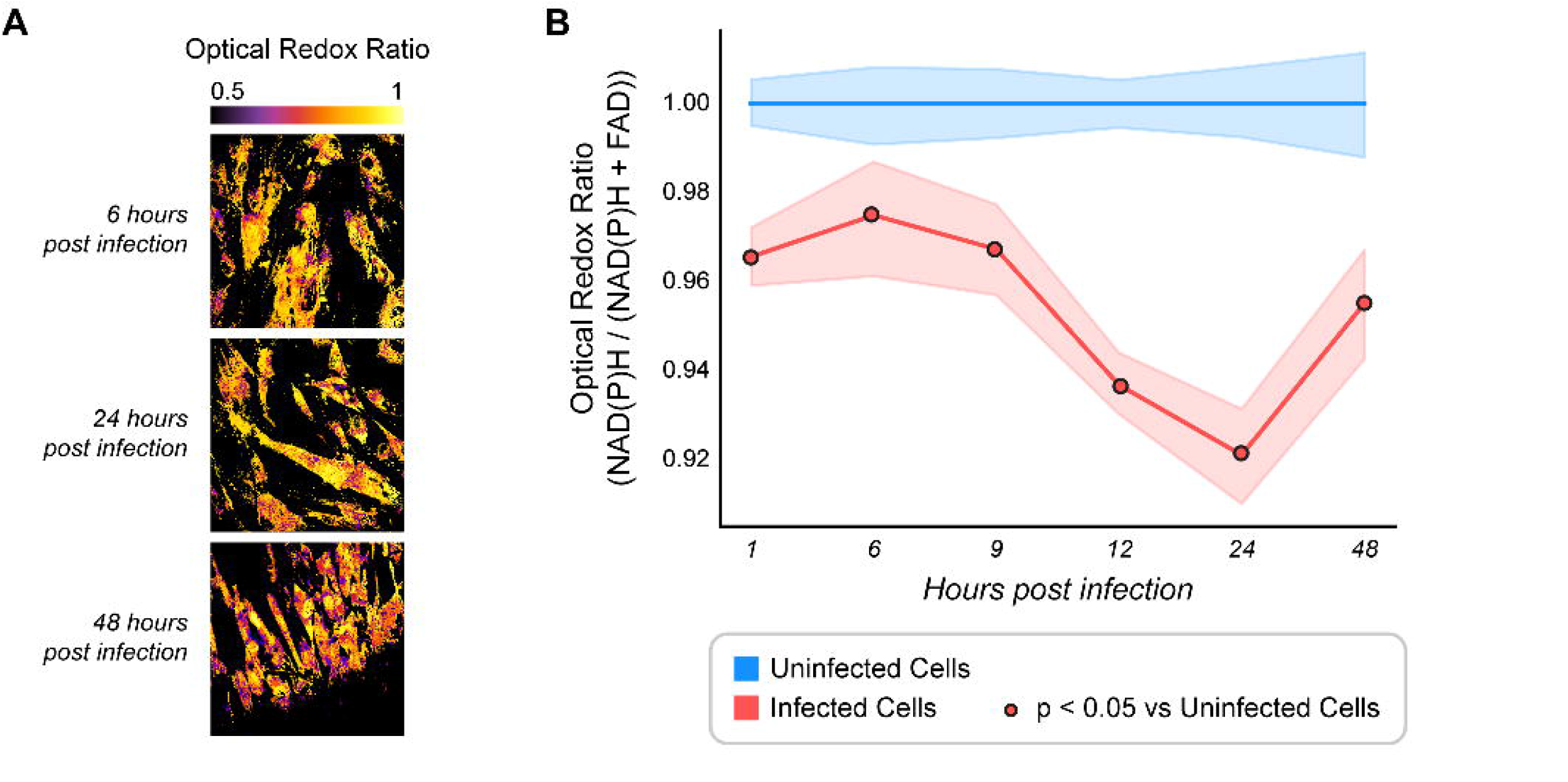
Optical Redox Ratio (ORR) temporal changes in *T. gondii* infected cells. **(A)** Representative images of temporal changes in optical redox ratio (ORR) of *T. gondii* infected HFF cells. Lower ORR means a more oxidized environment and higher ORR means a more reduced environment. Scale bar = 50 µm **(B)** ORR of *T. gondii* infected HFF cells vs. uninfected cells over a time course of infection, normalized to uninfected. Uninfected HFF cells are represented in blue and infected HFF cells are represented in red. Total number of cells n = 243, 225, 285, 277, 263, 312 for 1, 6, 9, 12, 24 and 48 HPI respectively. Each point represents the average of two independent experiments. Curves represent the mean and 95% confidence interval. Statistical significance was determined by an independent Student’s T-test. *p < 0.05. ps, picoseconds. Scale bar = 50 µm.

### NAD(P)H lifetime changed in uninfected vs T. gondii infected HFF cells

We obtained single cell qualitative (Figure 6A) and quantitative (Figure 6B) analysis of the mean NAD(P)H lifetime (τ_m_, Table 1) in uninfected HFF cells compared with high ME49 *T. gondii* infected cells. NAD(P)H τ_m_ increased in infected HFF cells at 1, 9, 12, and 24 HPI then decreased at 48 HPI compared to uninfected HFF cells (Figure 6B). Similarly, the proportion of protein-bound NAD(P)H (α_2_) in infected HFF cells increased at 1, 9, 12, 24, and 48 HPI compared to uninfected HFF cells (Figure 6C). NAD(P)H and FAD fluorescence of HFF cells changed differently with infection and reflect different metabolic activity during the 48-hour time-course in *T. gondii* infection (Figure 6, Figure S5B-D, Figure S6).

**Figure 6.**
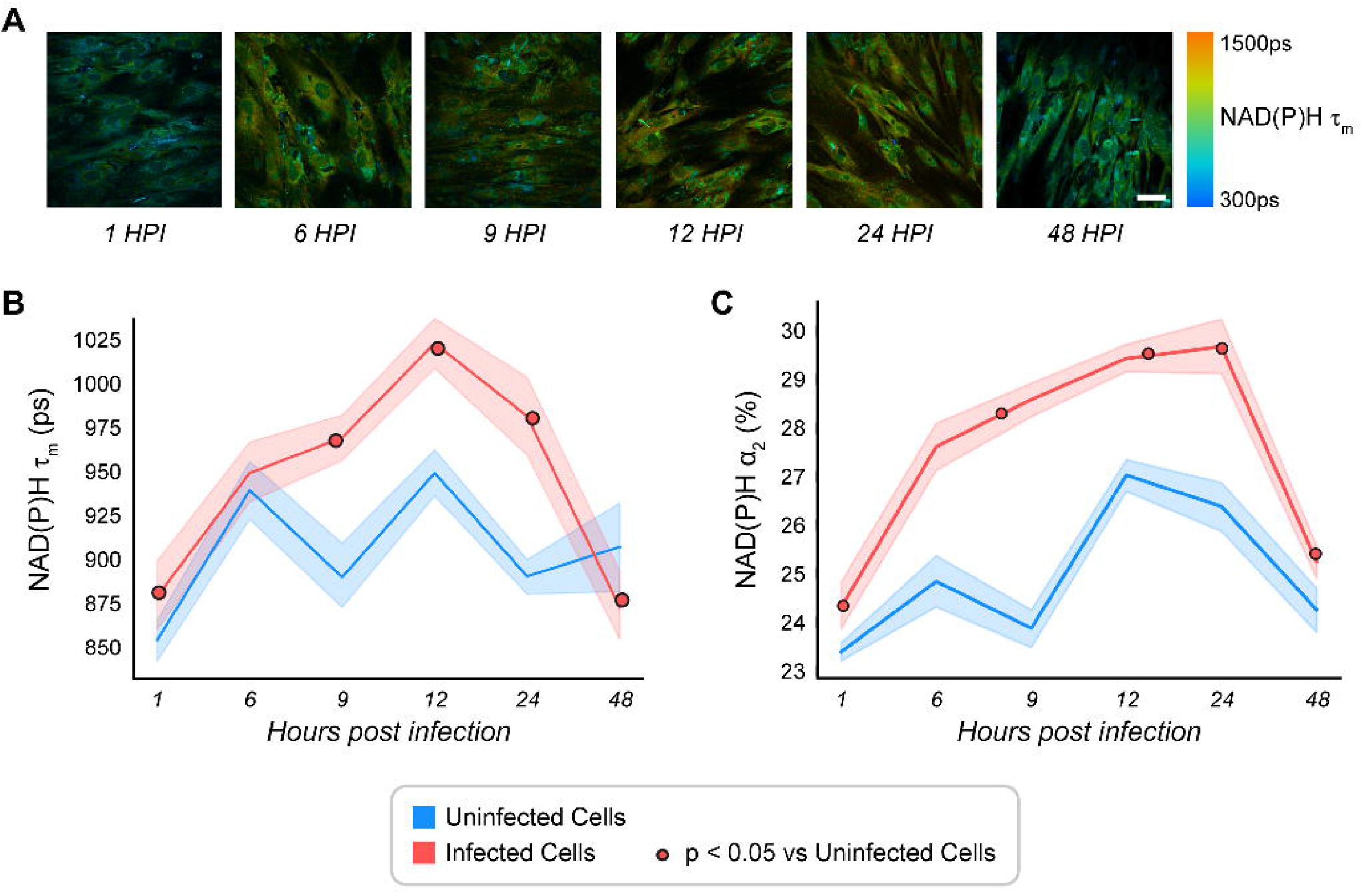
Temporal changes in NAD(P)H lifetime of *T. gondii* infected HFF cells. **(A)** Representative images of NAD(P)H mean lifetime (τ_m_) reported in picoseconds (ps) for 1 to 48 hours post infection (HPI). **(B)** NAD(P)H τ_m_ of *T. gondii* infected HFF cells in a 48-HPI time course experiment. **(C)** Percentage of protein bound NAD(P)H (α_2_) of *T. gondii* infected HFF cells in a 48 HPI time course experiment. Uninfected HFF cells are represented in blue and infected HFF cells (with *T. gondii* higher than 5%) are represented in red. Total number of cells n = 243, 225, 285, 277, 263, 312 for 1, 6, 9, 12, 24 and 48 HPI respectively. Each point represents the average of two independent experiments. Curves represent the mean and 95% confidence interval. Statistical significance was determined by an independent Student’s T-test. *p < 0.05. ps, picoseconds. Scale bar = 50 µm.

### T. gondii infection alters the host mitochondrial and glycolytic activity

We analyzed mitochondrial and glycolytic function to understand the mechanism of redox balance and NAD(P)H lifetime changes in *T. gondii* infection and validated our OMI results. These measurements have been standardized before with extracellular *T. gondii* [24] but not in infected cells or in intracellular parasites. We used a Seahorse XFp extracellular flux analyzer to measure the mitochondrial and glycolytic function of quiescent HFF host cells infected with wildtype *T. gondii* ME49 by a mitochondrial stress kit. As a control, we used a *T. gondii* type-I RH strain with and without a deletion in mitochondrial association factor 1 (RHΔMAF)[25].

Host mitochondria association (HMA) in *T. gondii* has been characterized previously and involves the *T. gondii* mitochondrial association factor (MAF) and 13 or more host proteins [25–28]. RH *T. gondii* strain infection on the host cells causes a relocalization of the host mitochondria around the parasite containing vacuole [28]. ME49 *T. gondii* strain does not have this phenotype or the MAF genes [25]. The cause and purpose of the mitochondria elongation in RH *T. gondii* infection is not totally clear. It could be a parasite strategy to induce host lipophagy, acquire fatty acids, amino acids, or pyruvate. It also could be a host defense mechanism that induces host mitochondria fusion that limits parasite proliferation [29].

We observed higher mitochondrial respiration measured in oxygen consumption rate (OCR) in the host cells infected with the parental *T. gondii* RH at 48 HPI as expected due to its association to host mitochondria (Figure 7A). HFF cells infected with *T. gondii* RHΔMAF showed reduced mitochondrial respiration. *T. gondii* ME49 infected HFF cells showed similar mitochondrial respiration as RHΔMAF infected HFF cells, probably due to the absence of host mitochondria association in both strains. Uninfected quiescent HFF cells showed low mitochondrial respiration as expected, suggesting that all changes observed correspond to the effect of intracellular parasite on host cells. We then compared the basal mitochondrial respiration in the four conditions (Figure 7B).

**Figure 7.**
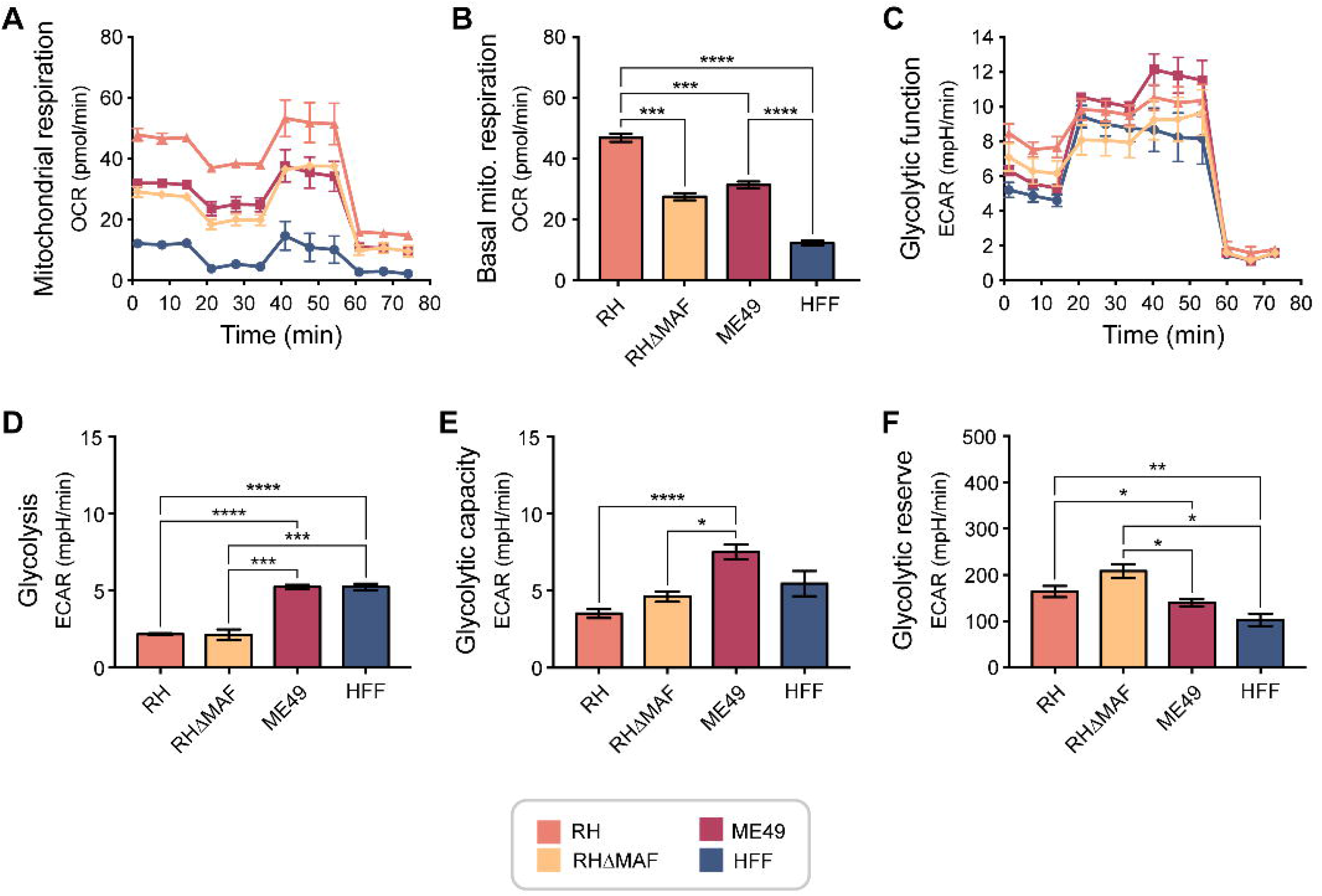
Mitochondrial and glycolytic activity in infected HFF cells using various *T. gondii* strains at 48 HPI. **(A)** Mitochondrial respiration **(B)** Basal mitochondria respiration. **(C)** Glycolytic function. **(D)** Glycolysis. **(E)** Glycolytic capacity. **(F)** Glycolytic reserve. *T. gondii* strains evaluated include: ME49, RHΔMAF, and RH. A and B were obtained by Seahorse mito stress kit and C-F were obtained by Seahorse glycolysis stress kit. All activities were measured by oxygen consumption rate (OCR) or by Extracellular acidification rate (ECAR). Each bar represents the mean of 6 or 12 replicates and error bars represent the SEM. Statistical analysis was performed by ANOVA, multiple comparisons were performed by Tukey’s test using Prism. *p < 0.05.

There is a significant increase in basal mitochondrial respiration in HFF infected with RH and ME49 compared to uninfected HFF cells (Figure 7B). Extracellular acidification rate (ECAR) was also measured to evaluate glycolytic activity with a glycolysis stress kit and seahorse analysis (Figure 7C – 7F). It was observed that ME49 infected cells have significantly more glycolysis at 48 HPI than RH strains (Figure 7C and 7D). Similarly, ME49 infected cells have significantly more glycolytic capacity at 48 HPI than RH strains (Figure 7E). However, ME49 infected cells have significantly less glycolytic reserve or less capability to respond to energetic demand at 48 HPI than RH strains (Figure 7F). Additionally, the infected HFF cells showed higher mitochondrial respiration than glycolytic function at 48 HPI, indicating that the parasite induces a metabolism predominantly based on mitochondria function in the infected host cell (Figure 7A and 7C).

We also performed Mitostress seahorse analysis in a time course experiment (Figure 8). We observed that RH and RHΔMAF infected HFF cells showed higher basal mitochondrial respiration than uninfected cells and ME49-infected HFF cells overtime (Figure 8A). ME49 infected HFF cells showed low basal mitochondrial respiration level and did not fluctuate over time (Figure 8A). RH infected HFF cells showed progressive increases in basal mitochondrial respiration levels until 36 HPI, then a reduction at 48 HPI (Figure 8A). This data suggests more mitochondrial activity in RH infected cells than ME49 infected cells in time-course infection. The Mitostress seahorse analysis also calculated ATP production in *T. gondii* infected HFF cells over time (Figure 8B). RH infected HFF cells and RHΔMAF infected HFF cells showed fluctuations in mitochondrial ATP during 48 HPI that were significant and not significantly, respectively (Figure 8B). ATP production in ME49 infected HFF cells did not fluctuate over time, like the uninfected cells (Figure 8B).

**Figure 8.**
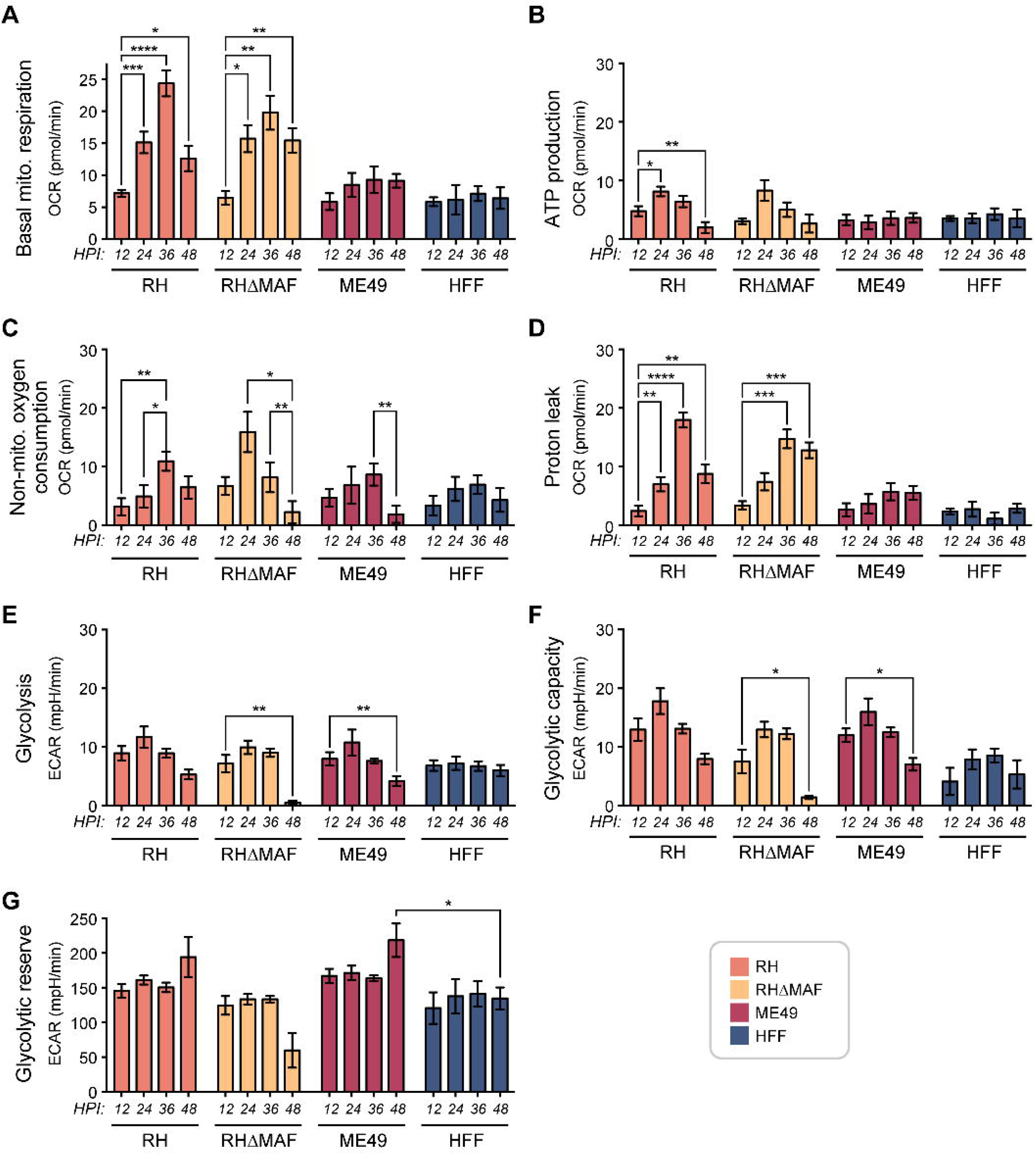
Temporal mitochondrial and glycolytic changes in HFF infected cells by different strains of *T. gondii* during a time course of infection. (A) Basal oxygen consumption rate. **(B)** ATP production. **(C)** Non-mitochondrial oxygen consumption. **(D)** Proton leak. **(E)** Glycolysis. **(F)** Glycolytic capacity. **(G)** Glycolytic Reserve. *T. gondii* strains evaluated include: ME49, RHΔMAF, and RH. OCR was calculated by Seahorse mito stress kit and ECAR was calculated by Seahorse glycolysis stress kit. Each bar represents the mean of 12 replicates and error bars represent the SEM. Statistical analysis was performed by two-way ANOVA, multiple comparisons were performed by Dunnett’s test using Prism. *p < 0.05.

Additional metabolic parameters evaluated included the non-mitochondrial oxygen consumption in HFF cells (Figure 8C). The non-mitochondrial oxygen consumption was calculated when the respiratory chain was totally inhibited with rotenone in combination with Antimycin; determining that most of the oxygen consumption is inhibited in the cell and the remaining oxygen consumption is attribute to non-mitochondrial oxidases and ROS production [30]. The peak of non-mitochondrial OCR was 36 HPI for RH and ME49 infected HFF cells and 24 hours for RHΔMAF infected HFF cells (Figure 8C). These results suggest that in infected HFF cells, other cytosolic mechanisms are consuming oxygen and producing energy, such as ROS production.

We also evaluated proton leak in the HFF cells. Proton leak increased progressively until 36 HPI, but then was reduced at 48 HPI in HFF infected with either of the two RH strains (Figure 8D). However, ME49-infected HFF cells showed lower proton leak than the RH infected HFF cells, probably due to factors such as less mitochondrial / ETC complexes damage; less passage of ions (as calcium or others) in inner membrane, or less electron slippage than RH strains (Figure 8D) [30]. Furthermore, we performed a glycolysis stress seahorse analysis in a time course experiment (Figure 8E-8G). We observed that the host cells infected with either of the three strains of *T. gondii*, fluctuated their glycolysis, glycolytic capacity, and glycolytic reserve during 48 hour-infection. Thus, the host cells infected with either of the three parasite strains have a fluctuation in glycolytic metabolism overtime, but the mitochondrial metabolism is predominantly active in RH infected cells.

### T. gondii infection alters the reactive oxygen species in the host cell

Oxidative stress results from an imbalance between the production of reactive oxygen species (ROS) and the ability of cells to scavenge them by the antioxidant system of the organism [31]. ROS include all highly reactive and unstable derivatives of molecular oxygen, such as hydrogen peroxide (H_2_O_2_), superoxide anion (O_2_) and the most dangerous hydroxyl radical (•OH). Because ROS is produced from oxygen metabolism, it is impossible to avoid ROS in aerobic organisms. They are generated in the cytosol and in organelles, such as mitochondria and peroxisomes. At physiological levels, ROS participate in cell signaling processes, but enhanced oxidative stress due to the excessive ROS formation may cause damage to all cellular macromolecules such as lipids, proteins, and nucleic acids, ultimately leading to cell death.

When *T. gondii* multiplies, it causes cellular disruption and cell death in an infected host. The resulting necrosis attracts inflammatory host cells, such as lymphocytes and monocytes. In the immune response against the parasite, large amounts of ROS are generated [32]. Oxidative stress resulting from the host response is toxic to parasites, but many studies have also reported that some consequences of parasitic infection in a host organism are the result of host defense mechanisms involving increased production of ROS [32]. We measured intracellular ROS in *T. gondii* infected HFF cells. ROS production fluctuated over time in infected HFF cells (Figure 9). The maximum ROS production was 24 HPI for ME49-infected cells and 48 HPI for RH infected cells (Figure 9).

**Figure 9.**
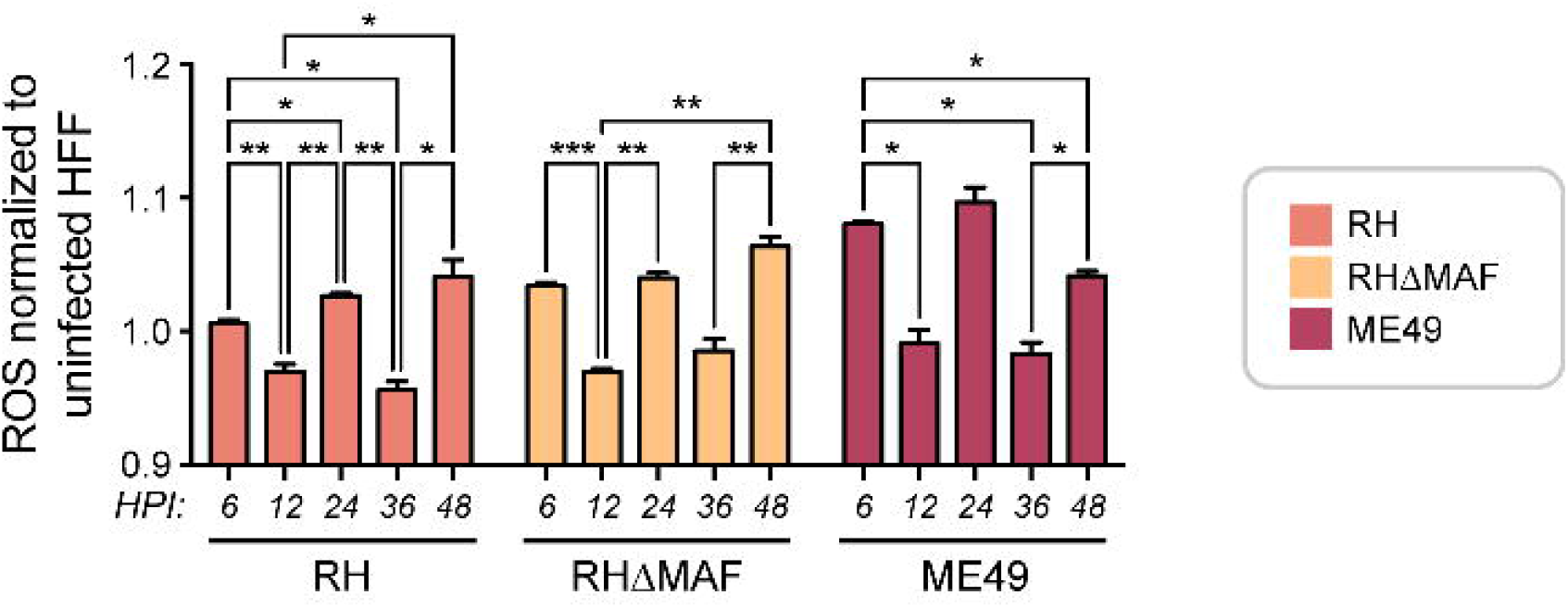
Reactive oxygen species (ROS) production in HFF infected with three different strains of *T. gondii* during a time course of infection. ME49, RHΔMAF, and RH *T. gondii* strains were evaluated. Each bar represents the mean of 4 replicates and error bars represent the SEM. Statistical analysis was performed by two-way ANOVA, mixed-effects model with Geisser-Greenhouse correction, Tukey multiple comparison test in Prism. *p < 0.05.

### T. gondii infection alters the host glucose and lactate production

As we observed a predominant glycolytic metabolism in *T. gondii* ME49 infected HFF cells (Figure 7A vs 7C), we analyzed intra- and extracellular glucose and lactate concentrations. Intracellular glucose concentrations revealed a progressive increase until 9 HPI, then it dropped significantly in ME49 *T. gondii* infected cells (Figure 10A). Intracellular lactate concentrations revealed a progressive increase until 6 HPI and then slightly started decreasing significantly in ME49 *T. gondii* infected cells (Figure 10B).

**Figure 10.**
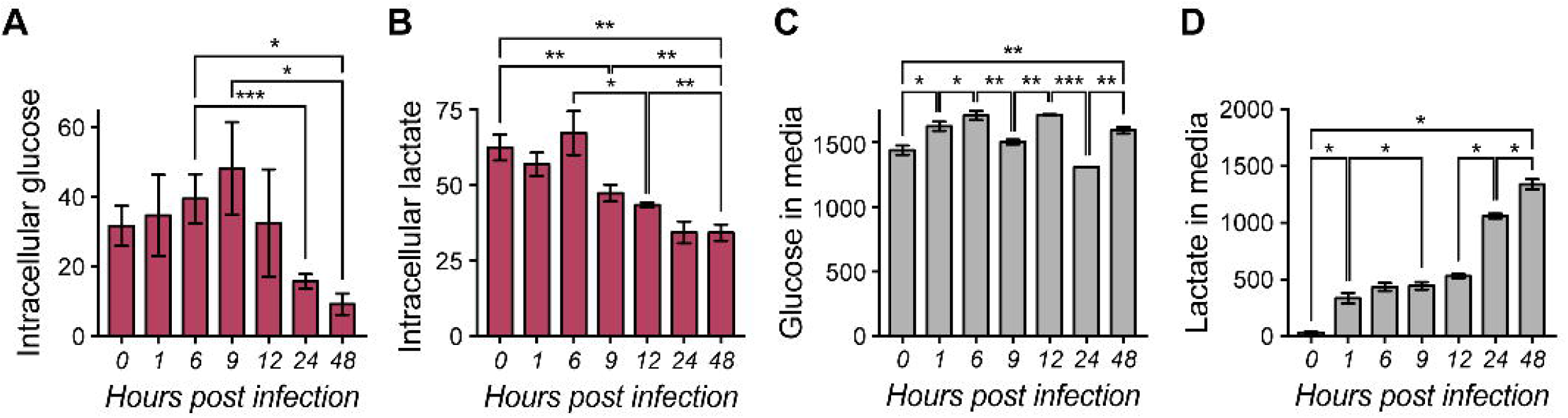
Glucose and lactate production in ME49 *T. gondii* infected host cells during a time course of infection. **(A)** Intracellular production of glucose in ME49 *T. gondii* infected HFF host cells during a time course infection, n= 6-15 wells. **(B)** Intracellular production of lactate in ME49 *T. gondii* infected HFF host cells during a time course infection, n= 6 wells. **(C)** Glucose in media in ME49 *T. gondii* infected HFF host cells during a time course infection, n= 4 wells. **(D)** Lactate in extracellular media in ME49 *T. gondii* infected HFF host cells during a time course infection, n= 6 wells. Each bar represents the mean of n replicates and error bars represent the SEM. pink = intracellular and blue = media / extracellular

These results correlate with the gene expression of enzymes involved in glycolysis during ME49 *T. gondii* infection to HFF cells, described in our previous publication (Figure S9) [5].The uptake of glucose from media significantly changed during 48 hour time course infection with more uptake at 9 and 24 HPI (Figure 10C). The concentration of lactate in extracellular media increased progressively and significantly during the 48-hour time course infection (Figure 10D). Infected cells switch to using lactate after depletion of glucose (24 and 48 HPI in Figure 10A and 10D), which may reflect the uptake and catabolism of this carbon source.

### T. gondii Kiss and Spit modify the optical redox ratio and NAD(P)H lifetime of the host cell

We performed an additional analysis called “kiss and spit” in which we used an actin polymerization inhibitor, cytochalasin D, which allows *T. gondii* to secrete the contents of their rhoptries into host cells while preventing infection [33]. Kiss and spit allow us to separate the effects of host manipulation due to secretion of rhoptries contents and the invasion. Cytochalasin D was administered during the different time-points of parasite infection by pretreating *T. gondii* with the inhibitor. Our previously published metabolomics study discovered that kiss and spit changes the host metabolism in nucleotide synthesis, the pentose, phosphate pathway, glycolysis, amino acid synthesis, and the abundance of the signaling molecules myo-inositol and cyclic-AMP [6]. Here, we demonstrated that kiss and spit alter many factors measurable by OMI. We found that kiss and spit reduced the ORR more than the normal infection (Figure 11A 11B, S7A, S7B, S8A and S8B). Additionally, kiss and spit increased NAD(P)H α_2_ (Figure 11C, S7C, S8C, S8D) and NAD(P)H τ_m_ (Figure 11D,S7D, S8E S8F) compared to cells treated with Cytochalasin D.

**Figure 11.**
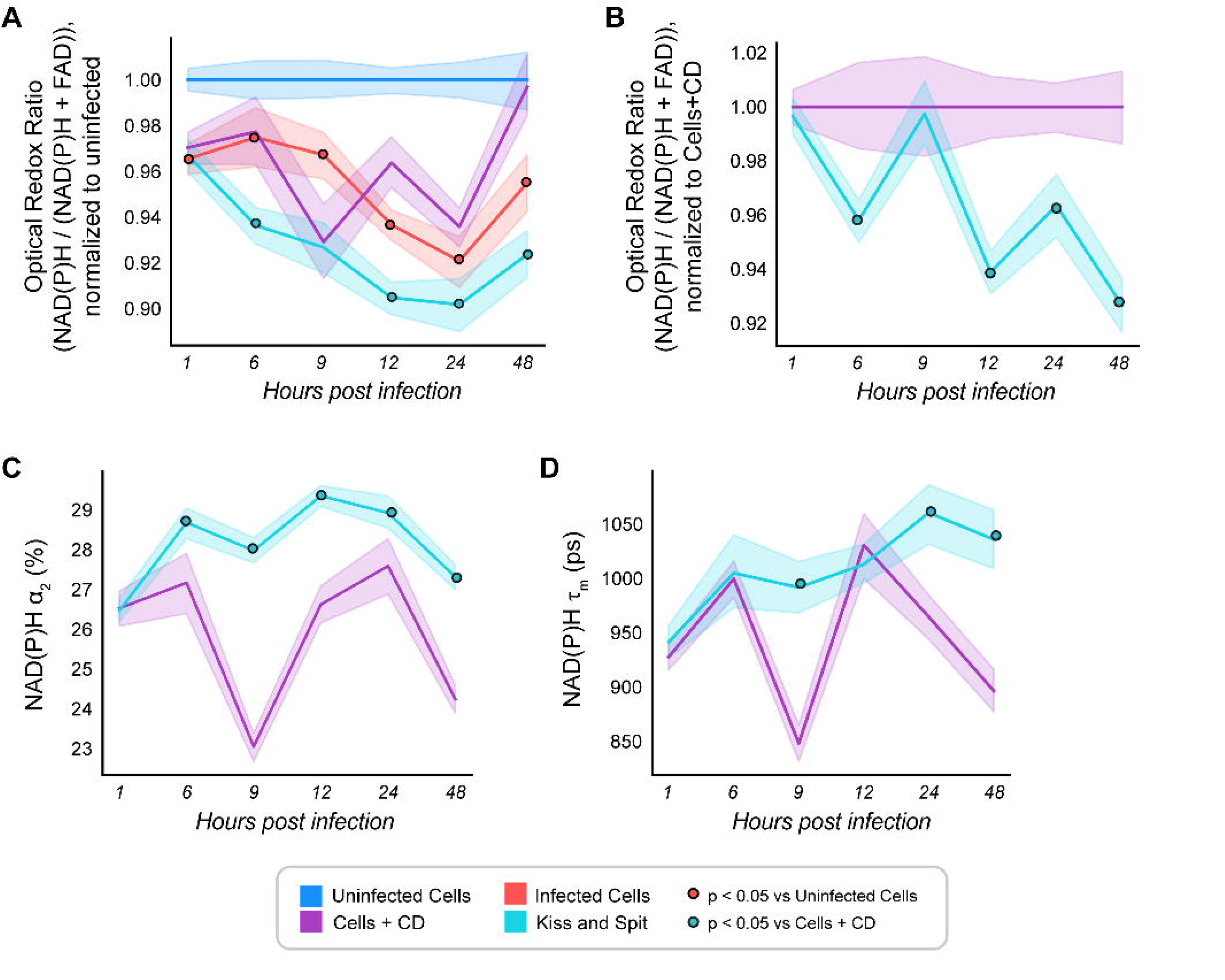
Temporal changes in OMI parameters during *T. gondii* kiss and spit. **(A)** Comparison of temporal Optical Redox Ratio changes in uninfected host cells, *T. gondii* infected host cells during active infection, kiss and spit, and in control of kiss and spit (Cell + Cytochalasin D (CD)). All groups normalized to uninfected cell control. **(B)** Temporal Optical Redox Ratio changes during *T. gondii* kiss and spit, normalized to its control (cells + CD). **(C)** Temporal changes in percentage of protein bound NAD(P)H (α_2_) during *T. gondii* kiss and spit. **(D)** Temporal changes in mean lifetime of NAD(P)H (τ_m_) during *T. gondii* kiss and spit. Uninfected HFF cells are represented in blue, infected HFF cells are represented in red, HFF cells treated with the inhibitor Cytochalasin D (CD) are represented in purple, and HFF cells with active *T. gondii* kiss and spit is represented in cyan. Each line represents the average of two independent experiments. Statistical significance was determined by an independent Student’s T-test. *p < 0.05. ps, picoseconds.

### NADP+ and NAD(P)H are associated to redox changes in ME49 T. gondii infection and kiss and spit

To determine the glucose contribution to the production of cofactors in *T. gondii* infection and kiss and spit, we performed a [^13^C] glucose labeling by LC/MS. We infected HFF with ME49 *T. gondii* and kiss and spit HFF cells at 9 HPI. Then, we labeled with [^13^C] glucose for 15 minutes and extracted the intracellular metabolites. We targeted the cofactors associated to redox changes: NAD+, NADP+, NADH, NADPH, and FAD. NAD+ (Figure 12A) and FAD (data not shown) also show some variations between infected vs non-infected cells and kiss and spit vs controls, but they were not significant. We found significant differences between infected and non-infected cells in NADP+ (Figure 12B), and NADH (Figure 12C). Similarly, kiss and spit produced a greater abundance of NADP+, NADH and NADPH compared to cells + cytochalasin D (Figure 12B, 12C,12D).

**Figure 12.**
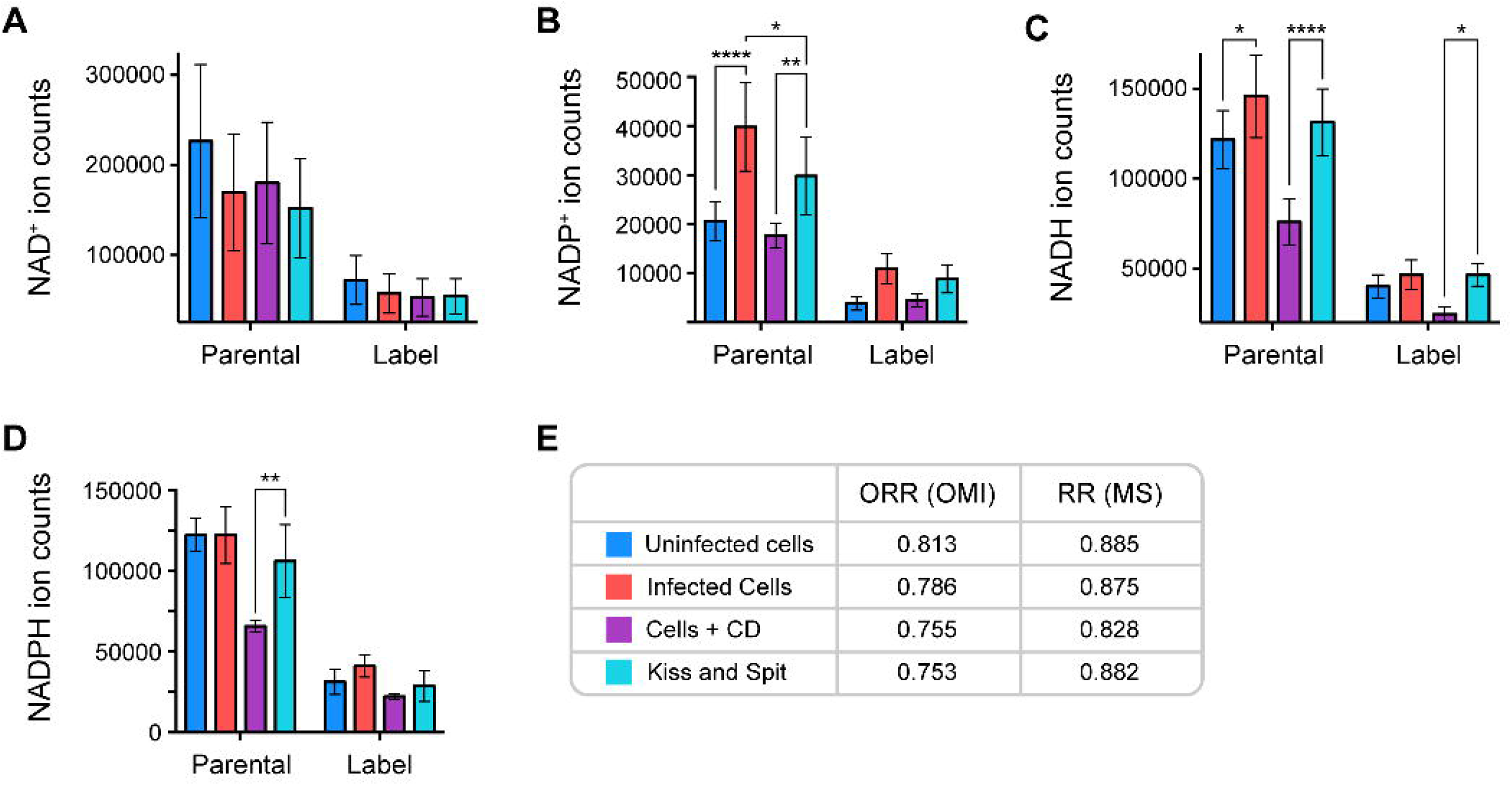
13C_6_ glucose labeling of intracellular NAD, NADP+, NADH and NADPH in *T. gondii* infection and Kiss and spit at 9 HPI by HPLC LC/MS. Ion counts of not-label (parental) and label (summatory of all label forms) of **(A).** NAD^+^. **(B).** NADP^+^. **(C).** NADH. **(D).** NADPH. Each bar represents the mean of two independent experiments with 7 replicates. Statistics were generated by a two-way ANOVA and Fisher LSD test. **(E).** Comparison of ORR calculated by OMI using NAD(P)H / (NAD(P)H + FAD formula, and RR (redox ratio) obtained by mass spectrometry using the formula (NADPH + NADH) / (NADPH + NADH + FAD) for the parental ion counts.

To evaluate whether there is a correlation between the optical redox ratio measured with OMI and the redox ratio measured with liquid chromatography/tandem mass spectrometry (LC/MS-MS), we compared both measures (Figure 12E). We used the average ORR calculated by NAD(P)H / (NAD(P)H +FAD) fluorescence at 9 HPI vs (NADPH + NADH) / (NADPH + NADH + FAD) calculated by LC/MS. Although there are differences between both techniques, we observed that the LC/MS redox ratio is similar to the ORR. Thus, infected cells showed a more oxidized redox ratio than uninfected cells (Figure 12E).

## DISCUSSION

Changes in host cell metabolism as a consequence of nutrient scavenging by intracellular parasites is difficult to study because of the inability to effectively separate parasite-derived activities from the host functions [4]. OMI is a non-invasive, label-free imaging technique that can be used to discover the features of intracellular parasite manipulation on host cell metabolism. This study brings insights into previously unexplored changes in host cell metabolism by *T. gondii* infection. OMI discovered shifts in host cell redox balance with *T. gondii* infection. Specifically, we found that ORR was more oxidized in ME49 *T. gondii* infected cells over time (Figure 5). These results indicate that the parasite manipulates the host cells at the redox biology level. Table 2 summarizes the factors that affect the optical redox ratio in ME49 *T. gondii* infected cells. It has been suggested that redox and ROS changes in host cells can influence *T. gondii* differentiation from tachyzoite to bradyzoite stage [34]. In our follow up studies, we will use FLIM to study the differences between *T. gondii* strains and their effect on host cell metabolism, the metabolic switch form tachyzoite to bradyzoite, and the different effects of *T. gondii* on different host cells, including immune cells.

**Table 2.** Factors that change Redox Ratio in ME49 *T. gondii* infected cells.

|  |  |
| --- | --- |
| ORR | Oxidized in infected cells |
| NAD(P)H Bound | Increase in Infected cells until 24 HPI |
| NAD(P)H Free | Decrease in Infected cells until 24 HPI |
| NAD(P)H $\tau_m$ | Increase in Infected cells until 24 HPI |
| Mitochondrial respiration | Does not change significantly during time course infection |
| Mitochondrial ATP production | Low. Does not change significantly during time course infection |
| Proton Leak | Low. Does not change significantly during time course infection |
| Non-mitochondrial Oxygen consumption | Increased progressively until 36 HPI, Collapsed at 48 HPI |
| Glycolysis | Increase until 24 HPI, then start reducing progressively. |
| Reactive Oxygen species (ROS) | High at 24 HPI. |
| Intracellular glucose | Increase until 9HPI, then reduce progressively |
| Extracellular Glucose | Fluctuate over infection |
| Intracellular Lactate | High production. Increase until 6 HPI, then reduce progressively |
| Extracellular Lactate | Increase over infection time. |

Similar to our results, previous studies found increased NADPH/NADP^+^ ratio associated with high oxidative phosphorylation activity in *T. gondii* infected myoblasts [34]. It was demonstrated by FLIM measurements that HCV induces increased NAD(P)H abundances in hepatic metabolism [18]. Oxidative pentose phosphate pathway (PPP) may be the main contributor of NAD(P)H in *T. gondii* parasite. Deletion of *T. gondii* Glucose-6-phosphate 1-dehydrogenase 2 (TgG6PDH2) enzyme, which is involved in the first step in the PPP, reduces the abundance of NADPH production, suggesting that this enzyme plays an important role in maintaining the cytosolic NADP/NADPH balance and tachyzoite anti-oxidant response [35]. We observed an increase in the amount of bound NAD(P)H with respect to free NAD(P)H in *T. gondii* infected HFF cells (Figure 6C and Figure S5C). Similar results were found in human cells infected with the intracellular bacteria *Chlamydia trachomatis* [19], where bound NAD(P)H was increased and free-NAD(P)H was reduced over the time course of infection measured by FLIM [19]. That study suggested that the reduction of free NAD(P)H is indicative of host cell starvation, by reduction of glycolysis. The reduction of free NAD(P)H was also observed in hepatic cells infected with HCV [18].

NADH and NADPH bind at least 334 known proteins in cells [36], and in infection [37], including enzymes important for diseases and *T. gondii* metabolism such as lactate dehydrogenase (LDH)[38], pyruvate dehydrogenase (PDH)[39], glucose 6-phosphate dehydrogenase (G6PDH)[40] and sirtuin 1 (SIRT1) [41]. The reason for increased bound NADPH (α_2_) in *T. gondii* infected host cells could be due to: (a) protein-bound NAD(P)H lifetime being sensitive to multiple fates of glucose carbon [36]; (b) the binding of NAD(P)H to different enzymes such as LDH, G6PDH or PDH, which could increase NAD(P)H α_2_ over time in *T. gondii* infected HFF cells (Figure 6C, Figures S10 - S12), or (c) host immune response mechanisms to eliminate the parasite [41].

In order to understand the molecular mechanism behind these changes observed by FLIM, we compared this results to our previously published RNA sequencing data on the same quiescent host cell system infected with ME49 *T. gondii* [5] and performed five different analysis. First, using the Database for Annotation, Visualization, and Integrated Discovery (DAVID), we analyzed the Go Terms related to molecular function of genes more abundant in ME49 *T. gondii* infected HFF cells (Supplemental Figure S10). In all time points, we observed increased abundance of Go terms related to NADH-dehydrogenases, hydrogen ion transmembrane transport, proton-transporting ATPases, electron carrier, Cytochrome C oxidase and oxidoreductase activity. These NADH dehydrogenase enzymes could be binding to NADPH (Figure 6C). Second, we analyzed the reactome of genes upregulated in ME49 *T. gondii* infected HFF cells (Figure S11). In all time points, we observed an increased abundance of genes clustered in different categories such as metabolism, respiratory electron transport, TCA cycle, cellular response to stress, ROS and reactive nitrogen species (RNS) production, and mitochondrial biogenesis. These genes could be upregulated in response to the *T. gondii* infection and affect the redox balance of host cell. Third, we analyzed the most abundant NAD(P)H-dependent enzymes (Figure S12)[5]. We assume that because there are more than 370 NAD(P)H-dependent enzymes, only the most abundant enzymes will have an impact on the resulting fluorescence lifetime of NAD(P)H and will preferentially bind to the coenzymes [37]. Time-course gene expression shows the most likely host enzymes to bind to NAD(P)H during *T. gondii* infection (Figure S12) similar to other parasites using FLIM [37]. We want to highlight the abundance of lactate dehydrogenase (LDH), Phosphoglycerate dehydrogenase (PHGDH), Succinate dehydrogenase subunit D, Nitric oxide synthase (NOS2), Isocitrate Dehydrogenase 3a, and Dual oxidase 1 (DUOX1), among others (Figure S12). Fourth, we also analyzed *T. gondii* gene expression of 58 genes related to redox biology [42] (Figure S13). At 48 HPI, the most expressed enzymes are *T. gondii* superoxide dismutase, catalase, and proteins with Thioredoxin domains (Figure S13). Fifth, Figure S14 shows the abundance of other genes that should also be related to redox biology in *T. gondii* metabolism. We did not identify the host NADPH–oxidase complex (NOX), another important bound or dependent NAD(P)H enzyme that plays an important role in *T. gondii* infection [43–45].

Previous studies have demonstrated that changes in ORR and NAD(P)H lifetimes correlate with changes in mitochondrial and glycolysis, measured by seahorse analysis [9,46–48]. Using this technique, we found that host cells infected with *T. gondii* RH parental strain, which has the MAF factor, show high mitochondrial respiration, more ATP demand (Figures 7 and 8), altered oxidative phosphorylation and/or mitochondrial gene expression reported previously, probably related to its association to host mitochondria [25,29,49]. *T. gondii* ME49 showed similar mitochondrial respiration to RHΔMAF (Figures 7 and 8), probably due to the absence of host mitochondria association in both strains [30]. ME49 infected cells do not produce mitochondrial ATP equal to RH strains and it does not fluctuate over time (Figure 8B) but have abundance in glycolysis (Figure 7D and 7E). Prior studies indicate that *T. gondii* is capable of maintaining cellular ATP homeostasis via either glycolysis or mitochondrial oxidative phosphorylation [50]. These results suggest that ORR and NAD(P)H changes observed by FLIM correspond to alterations in the glycolytic metabolism of ME49 infected cells.

There are other cytosolic mechanisms that consume oxygen and affect the redox biology of the cell [30]. For this reason, we measured ROS production in infected cells. Our results showed that the infected cell ROS production fluctuated over time (Figure 9). It has been demonstrated previously that host ROS production can inhibit *T. gondii* growth [51]. Human patients infected with *T. gondii* showed increase oxidative stress markers and reduction of antioxidant markers [52]. Elevated ROS levels have been previously reported in *T. gondii* infected host cells [34,43,53]. Asymptomatic *T. gondii* seropositive cats also showed an increase in ROS levels [54]. The rapid release of ROS plays a significant role against *T. gondii*, but also contributes to oxidative injury inflicting tissue damage and disease pathology. ROS production in *T. gondii* infection induces host DNA damage [53] and intracellular lysosomal membrane damage, which is followed by apoptosis or necrosis [31]. These results suggest that ROS production in ME49 infected cells would be an important factor affecting the ORR and NAD(P)H changes found by FLIM.

To understand the molecular mechanism behind the NAD(P)H and redox changes observed by FLIM, we analyzed glucose and lactate fate and flow during *T. gondii* infection. Glucose is the preferred nutrient for *T. gondii* and its assimilation via glycolysis supports the optimal growth of the parasite [50,55]. Here, we found a reduction of intracellular glucose in *T. gondii* infected host cells and an increase of lactate exportation over time (Figure 10). This reduction in host intracellular glucose, from 9 HPI and forward, correlates with the oxidized ORR quantified by OMI (Figure 5 and 10A). Similar results were observed in myoblasts and myotubes infected with *T. gondii* type II [34]. Many studies have demonstrated that *T. gondii* can propagate in the absence of glucose, using glutamine or acetate as an alternative source of energy [50,56]. Our intracellular measurements of glucose and lactate correlate with the intracellular metabolites [5] and up-regulation of glycolytic enzyme gene expression [4,5] (Figure S9). *T. gondii* and host lactate dehydrogenase (LDH) activity is up-regulated during the infection (Figure S9J) [4,5]. In *T. gondii* infected host cells, lactate exportation was observed to progressively increase (Figure 10D). Pyruvate and lactate metabolites are abundant in *T. gondii* infection because the parasite must maintain a pyruvate homeostasis [57]. Both metabolites serve as a circulating redox buffer that equilibrates the NADH/NAD^+^ ratio in cells [58]. Lactate serves as a major circulating carbohydrate fuel [58], is exported from the parasite [59], and helps to regulate the redox balance in infection, cancer, and immune cells [60–62]. Lastly, lactate is responsible for 30% inhibition of *T. gondii* tachyzoite to bradyzoite conversion in Vero cells [63]. It has been demonstrated that low glucose and high lactate environments are immunosuppressive; these conditions are found in the placenta, gastrointestinal tract, and in tumor microenvironment [60], as well as *T. gondii* infection (Figure 10).

To understand the biological relevance of an optically derived redox ratio and the metabolic pathways that contribute to it in *T. gondii* infection, we performed a [^13^C] glucose flux analysis (Figure 12). Redox has been associated with intracellular concentrations of NADH and NAD^+^ in stem cells [64]. Previous studies found no production of NAD+ in *T. gondii* [65], and our data did not find significant differences in NAD+ levels with infection (Figure 12A). The changes in ORR (Figure 5) and relative amounts of bound and free NAD(P)H (Figure 6C) observed in *T. gondii* infection by OMI correspond to NADH or NADPH, the autofluorescence cofactors, which showed significant differences with infection by LC/MS (Figure 12C and 12D) [66]. LCM/MS found a significant abundance of NADP+ in ME49 *T. gondii* infected cells and kiss spit (Figure 12 B). The production of NADP+ is important for *T. gondii* because isocitrate dehydrogenase, the rate limiting step of the TCA cycle, is NADP+ dependent, not NAD+ dependent as the human enzyme [5,65]. Uninfected vs Infected cells ORR showed a similar trend as the biological redox ratio obtained by metabolomics (Figure 12E).

Finally, we determined by OMI how *T. gondii* rhoptry contents discharged during kiss and spit remodel the HFF metabolism. Kiss and spit is a pre-invasion process where the contents of the *T. gondii* rhoptry organelles are secreted into the host cytoplasm, even in uninfected cells [67,68]. The rhoptries contain an estimated fifty proteins and lipids, most of which are functionally uncharacterized, [68–73]. Cytochalasin D or Mycalolide B function as actin polymerization inhibitors that prevent invasion, allowing the host changes associated with kiss and spit to be studied independently of parasite invasion and replication [33,74–77]. We found that kiss and spit reduced the ORR (Figure 11, S7) and the free NAD(P)H (α_1_) (data not shown) in a similar trend to normal infection (Figure S8). Kiss and spit increased the relative amount of bound NAD(P)H (α_2_) and the NAD(P)H mean lifetime (Figure S7) as the full infection does. This finding suggests that the *T. gondii* kiss and spit manipulates the redox biology of the host cells similar to full infection, as has been demonstrated before using several reporter systems [78].

## Conclusion

OMI has been used to understand the metabolic changes in numerous disease models. We investigated whether OMI could be used in *T. gondii* infection to determine the changes induced in the host cell by the parasite and evaluated the effect of infection on redox balance in the host cell. Our results concluded that *T. gondii* infected HFF cells show a more oxidized ORR than the uninfected cells. Many host and parasite genes (described in Figure S11-S15), could be related to these changes in redox biology, especially the ones implicated in ROS production (Figure 9). *T. gondii* infected HFF cells showed an increase in the relative amount of bound NAD(P)H with respect to free NAD(P)H over a time course of infection. This abundance of bound NAD(P)H in *T. gondii* infected HFF cells could be correlated to the abundance of enzymes that bind to NAD(P)H with *T. gondii* infection such as dehydrogenases, LDH, PHGDH, GAPDH, DUOX1 (Figure S13), or to the abundance of PPP and the high rate of glycolysis in the intracellular ME49 *T. gondii* parasite. Particularly, this strain of parasite depends more on glycolysis, and less on oxidative phosphorylation (Figure 7 and 8); since it does not have the mitochondrial association factor as the RH strains have. Finally, we explored changes associated with kiss and spit using OMI. Kiss and spit also showed a more oxidized ORR in the host cell and increases in the relative amount of bound NAD(P)H with respect to free NAD(P)H. Separate analysis of host and pathogen metabolism is still challenging and requires novel experimental and technical approaches that facilitate dynamic monitoring of metabolic changes inside the intracellular parasite separately from the host cell in living cells. A future OMI analysis of *T. gondii* parasite requires magnification of the images and segmentation of the intracellular and extracellular parasite.

A second limitation of NAD(P)H autofluorescence imaging is that conversion of NAD(P)H fluorescence intensity values to absolute concentration values is not straight forward because the different quantum yields of free and protein-bound NAD(P)H have to be calculated [19,79,80]. The fluorescence decay parameters of the phosphorylated and non-phosphorylated forms of reduced NAD are the same and are indistinguishable. Although estimations of cellular concentrations suggest that a substantial part of the cellular fluorescence originates from NADPH rather than from NADH [81], we and others have demonstrated significant NAD(P)H fluorescence lifetime changes by inhibiting glucose metabolism corresponding to projected changes of cellular NADH concentration [79,82,83].

A more detailed understanding of the metabolic activity and needs of *T. gondii* during the intracellular growth phase is needed to conceive novel therapeutic strategies that target the pathogen in its intracellular growth phase without affecting the host. More broadly, fluorescence lifetime imaging using two-photon microscopy reveals new insights into the crosstalk between host and pathogen metabolism and suggests manipulating *T. gondii*-induced changes in subcellular NAD(P)H contents and redox biology. In the process of understanding how intracellular pathogens interfere with host cell metabolism, metabolic profiling of infected cells by OMI will be an invaluable tool that complements established large scale genomic and proteomic approaches.

## MATERIALS & METHODS

### *T. gondii* Strains and Cell Culture

Low passage mCherry type II-ME49 *T. gondii* was used for OMI. Low passage type II ME49 *T. gondii* was used for the rest of the experiments. The parental strain RHΔKU80 (RH) and the modified RHΔKU80ΔMAF (RHΔMAF) strains obtained from Dr. Boothroyd were used as controls for Seahorse and ROS analysis. Human Foreskin Fibroblasts (HFFs) were grown in DMEM with 10% Fetal Bovine Serum (FBS), 2 mM L-glutamine, and 1% penicillin-streptomycin (Sigma-Aldrich). Once HFFs cells were in deep quiescence, defined as 10 days post confluency, DMEM media was changed to metabolomic media for all metabolomic analysis. Metabolomic media is made of RPMI1640 supplemented with 2 mM L-glutamine, 1% FBS dialyzed against PBS (MW cutoff of 10 kD), 10mM HEPES, and 1% penicillin-streptomycin.

### Time Course Infection and Kiss and Spit

For OMI, HFF dishes in metabolic media and in triplicated were treated as follow: (a) Uninfected; (b) infected with 2 x 10^6^ ME49 *T. gondii* tachyzoites; (c) infected with 2 x 10^6^ ME49 tachyzoites that had been pre-incubated with 1.5 μM cytochalasin D during 15 minutes at 37°C (Sigma-Aldrich); (d) incubated with 1.5 μM cytochalasin D (Sigma-Aldrich). Cytochalasin D was kept during all the length of experiments and imaging. At time points 1, 6, 9, 12, 24 and 48-HPI, dishes were imaged and maintained during imaging at 37°C and 5% CO_2_ using a stage-top incubator system (Tokai Hit).

### Metabolomics

HFF dishes in triplicate were treated as the four conditions described previously. At time point 9-HPI, dishes were washed three times with ice cold PBS, and incubated for 15 minutes with metabolomic media plus Glucose-^13^C_6_ (sigma) (1g/L). Then, the dishes were quenched with 80:20 HPLC grade Methanol: Water (Sigma-Aldrich).

Dishes were incubated on dry ice at −80°C for 15 minutes. Plates were scraped, the solution removed, and spun at 2500 x g for 5 minutes at 4°C. The supernatant was removed and stored on ice, then the pellet was washed again in quenching solution and re-spun. Supernatants were combined, dried down under N_2_, and stored at −80°C. Samples were resuspended in 100 µL HPLC grade water (Fisher Optima) for analysis on a Thermo-Fisher Vanquish Horizon UHPLC coupled to an electrospray ionization source (HESI) part of a hybrid quadrupole-Orbitrap high resolution mass spectrometer (Q Exactive Orbitrap; Thermo Scientific). Chromatography was performed using a 100 mm x 2.1 mm x 1.7 µm BEH C18 column (Acquity) at 30°C. 20 µL of the sample was injected via an autosampler at 4°C and flow rate was 200 µL/min. Solvent A was 97:3 water/methanol with 10 mM tributylamine (TBA) (Sigma-Aldrich) adjusted to a pH of 8.2 using approximately 9 mM Acetate (final concentration, Sigma-Aldrich). Solvent B was 100% methanol with no TBA (Sigma-Aldrich). Products were eluted in 95% A / 5% B for 2.5 minutes, then a gradient of 95% A / 5% B to 5% A / 95% B over 14.5 minutes, then held for an additional 2.5 minutes at 5%A / 95%B. Finally, the gradient was returned to 95% A / 5% B over 0.5 minutes and held for 5 minutes to re-equilibrate the column. MS parameters included: scan in negative mode; scan range = 70 - 1000 m/z; Automatic Gain control (AGC) = 1e6, spray voltage = 3.0 kV, maximum ion collection time = 40 ms, and capillary temperature = 350C. Peaks were matched to known standards of NADH, NAD^+^, NADPH and NADP^+^ for identification. Data analysis was performed using the Metabolomics Analysis and Visualization Engine (MAVEN) software.

### Two-Photon Imaging

Fluorescence lifetime images were taken on a custom-built inverted multiphoton microscope (Bruker Fluorescence Microscopy, Middleton, WI, USA), as previously described [84–86]. Briefly, the system consists of an ultrafast laser (Spectra Physics, Insight DS-Dual, Milpitas, CA, USA), an inverted microscope (Nikon, Eclipse Ti, Tokyo, Japan), and a 40*×* water immersion (1.15NA, Nikon) objective. NAD(P)H and FAD images were acquired sequentially for the same field of view using an excitation wavelength of 750 nm and a 440/80 nm emission bandpass filter for NAD(P)H fluorescence, and an excitation wavelength of 890 nm and a 550/100 nm emission bandpass filter for FAD fluorescence. mCherry *T. gondii* was excited at 1090 nm with emission at 690/50 nm. During imaging, dishes were maintained at 37°C and 5% CO_2_ using a stage-top incubator system (Tokai Hit). Fluorescence lifetime images were collected using time-correlated single photon counting electronics (SPC-150, Becker and Hickl, Berlin, Germany) and a GaAsP photomultiplier tube (H7422P-40, Hamamatsu Photonics, Hamamatsu, Japan). A pixel dwell time of 4.8 µs was used to acquire 256 *×* 256-pixel images over 60 s total integration time. The photon count rates were maintained at 1–2 *×* 10^5^ photons/second to ensure adequate photon observations for lifetime decay fits, and no photobleaching. The instrument response function was measured from second harmonic generation of urea crystals excited at 900 nm, and the full width at half maximum (FWHM) was calculated to be 220 ps.

### Quantification of Fluorescence Lifetime Components

NAD(P)H and FAD fluorescence lifetime images were analyzed using SPCImage software (Becker & Hickl, Berlin, Germany) as previously described [87]. At each pixel, the fluorescence lifetime decay curve was deconvolved with the instrument response function and fit to a two-component exponential decay model, I(τ) = α_1_ *×* exp(*−*τ/ τ_1_) + α_2_*×* exp(*−*τ/τ_2_) + C. In this model, I(τ) is the fluorescence intensity at time *t* after the laser excitation pulse, *α* represents the fractional contribution from each component, *C* accounts for background light, and t represents the fluorescence lifetime of each component [15,87]. A two-component model was used because both NAD(P)H and FAD can exist in two conformational states, bound or unbound [17,88]. We use a binning factor of 1 or 2.

### Creating HFF whole cell masks

Initial whole cell masks were created manually using the software CellProfiler. These masks were then further revised and validated for accuracy by a scientist using the Napari image viewer, by overlaying the mask over the NAD(P)H intensity image and improving the cell boundaries and/or labeling regions missed by CellProfiler. Whole cell was defined as the cell border including the nuclei.

### Calculating Optical Redox Ratio, NAD(P)H and FAD parameters

The total NAD(P)H and FAD intensity was calculated from the lifetime data by summing the number of photons detected for every pixel in the cell mask. The intensity of NAD(P)H was then divided by the sum of FAD plus NAD(P)H intensity for each pixel to calculate the optical redox ratio (Table 1). Values for τ_1_, τ_2_, α_1,_ α_2_ of NAD(P)H and FAD were measured for each host cell too (Table 1). The mean lifetime (τ _m_) of both NAD(P)H and FAD were calculated as τ_m_ = α_1_ τ_1_ + α_2_ τ_2_. With the whole cell masks, the *T. gondii* masks and the SPCImage exports (τ_1_, τ_2_, α_1_, α_2_ for NAD(P)H and FAD), we extracted the OMI features for each cell for all the images collected. In python, we loaded all the images as sets containing corresponding masks and images. We then used the function *regionprops_omi* within the *cell-analysis-tools* [89] python library to extract features for each image for each cell and saved the output as a CSV file. All these resulting files were then aggregated into a single CSV file and used for data analysis and to generate figures. Statistical significance was added to the figures by the statannotations python package [90] using an independent t-test.

### Quantifying *Toxoplasma gondii* in mCherry images

In python, the mCherry intensity image was loaded and the intensity of the top 5% of the pixels was mapped to equal the 95 percentile intensity value, then from the resulting image the top 10% brightest pixels of the image were kept (Figure 1). The resulting image was then binarized and then dilated using an octagon with footprint (1,1). Binarization was done by making all pixels with values greater than 0 equal to 1. Holes in the connected components were filled using the *binary_fill_holes* function from the image processing library *scikit-image* [91]. Followed by the function *remove_small_objects* from the same library to remove small sections of connected components less than 30 pixels in area. Another version of the mask was then created by taking the original NAD(P)H intensity image and keeping the 5% brightest pixels. Lastly, these two images were combined using a bitwise OR operation to create the final binary mask (Figure 1).

### Quantifying intracellular *Toxoplasma* and establishing a 5% threshold

After creating the whole cell masks and the *T. gondii* masks for the datasets, both masks were loaded into python to quantify the amount of *T. gondii* in each cell according to their overlap as shown in Figure 1. This was done by multiplying the final *T. gondii* mask with the final host cell mask to produce a mask capturing the amount of parasite in each cell. This new mask has the same region label for the *T. gondii* pixels as the pixels in the whole cell mask. We then divide the sum of the pixels in the mask containing *T. gondii* content inside the cell by the number of pixels in the whole cell area for every cell (Figure 2A). This quantified the percent of *T. gondii* infection in each cell. *T. gondii* did not infect all cells equally; to compare the percentage of infection, we plotted a histogram of percent *T. gondii* infection by cells for each timepoint, for each experiment, and determined empirically that the lower 5% had no significant *T. gondii* infection or the infection could be a false positive due to pixel noise captured by the *T. gondii* masks.

### Metabolic Profiling-Seahorse analyzer

HFF cells were seeded in a seahorse 96 well plate and allow them to reach confluency and quiescent for two weeks. After this time, all steps were performed in metabolic media. Then, cells were infected with 6,000 taquizoites / well from the three different strains of *T. gondii*: ME49, RHΔMAF and RHΔKU80. The control group was the HFF cells without infection. After 48 HPI (Figure 7) or during time-course infection (Figure 8), Mito stress analysis was performed using 1 µM Oligomycin, 2 µM Carbonyl cyanide-4 phenylhydrazone (FCCP) and 0.5 µM rotenone. Glycolysis stress analysis was performed using 10 mM Glucose, 2 µM Oligomycin and 50 Mm of 2-DG. Seahorse analysis was performed in a Seahorse Bioscience XF96 Extracellular Flux Metabolic Analyzer in the Small Molecule Screening Facility at the University of Wisconsin.

Oxygen consumption rate (OCR) and extracellular acidification rate (ECAR) was calculated [9]. Mitochondrial respiration is measured by OCR and is a quantitative metric of mitochondrial function via oxidative phosphorylation (OXPHOS). Glycolysis is indicated by ECAR [92]. Graphs and statistical analysis were performed in Wave software v 2.6.3.5 from Agilent technologies and statistical analysis was performed in prism.

### ROS labeling

Live infected and non-infected HFF cells were stained with 5 μM CellROX™ Green Reagent in complete medium for 30 minutes at 37°C, 5% CO_2_. Fluorescence was measured in an Incucyte machine. Reactive oxygen species (ROS) were labeled using the CellROX™ Green Reagent according to the manufacturer instructions (Molecular Probes, Eugene, USA). CellROX™ Green predominantly detects hydroxyl radicals and superoxide anions and only to low extent tert-butyl-hydroperoxide and do not detect hydrogen peroxide (www.thermofisher.com)[34,93].

### Lactate and glucose assays

Quiescent HFF cells in a 24-well plate were infected with 2×10^5^ ME49 *T. gondii* tachyzoites per well in cell metabolic culture media. Extracellular and intracellular samples were collected from three biological replicates for each of the seven time-points after *T. gondii* infection time-course. For lactate and glucose intracellular assays samples were homogenized by sonication. Abcam Glucose assay kit (ab65333) and Abcam Lactate kit (ab65330) were used. Samples were read in a plate reader at 533/587 nm. Concentrations in pMol were determined by standard curve with glucose and lactate standards.

### Gene expression analysis

We re-analyzed our previous published gene expression analysis (accession number PRJNA497277)[5]. This data corresponds to ME49 *T. gondii* infected HFF cells. Lists of significant genes were used as input for gene ontology enrichment analysis using the Database for Annotation, Visualization, and Integrated Discovery (DAVID, v6.8). First, we performed clustering by Go term of significant genes during each time point. Second, we performed clustering by significant genes related to reactome during each time point. Third, we generated heatmaps of time-course gene expression of the most likely host enzymes that bind to NAD(P)H during *T. gondii* infection, similarly as it has been published previously for other parasites using FLIM [37]. Fourth, we analyzed the expression of 58 genes related to redox biology in *T. gondii.* Fifth, we analyzed the expression of other genes related to redox biology in *T. gondii*.

### Statistical analyses

Biological repeats are defined as separate time-point infections collected on separate days. Statistical significance was set to 0.05. Graphical displays were generated in Python using the open-source graphing package Matplotlib (https://matplotlib.org/) and Seaborn (https://seaborn.pydata.org/). Each data point is a different cell; boxplots show median (central line). Statistical significance was added to the figures by the statannotations python package [90] using an independent t-test. Prism was used to create graphs and run statistical analysis.

## Supplementary Figures

**Figure S1.** Percentage of intracellular *T. gondii* infection per cell during time course infection in two independent experiments. (**A)** Experiment #1. **(B)** Experiment #2. X-axis represents the percentage of intracellular parasite by cell area by time point using 5% bin sizes. There are six time points: 1, 6, 9, 12, 24, and 48 HPI. The Y-axis represents the cell count, number of cells in each specific bin.

**Figure S2. Percent of intracellular *Toxoplasma gondii* infection per cell during time course infection in experiment #1.** There are six time points: 1, 6, 9, 12, 24, and 48 HPI. Each time point is represented by one graph. X-axis represents the percentage of intracellular parasites. The Y-axis represents the cell count, number of cells in each specific percentage. **(A)** Time point 1 HPI = 113 cells. **(B)** Time point 6 HPI = 150 cells. **(C)** Time point 9 HPI = 187 cells. **(D)** Time point 12 HPI = 196 cells. **(E)** Time point 24 HPI = 187 cells. **(F)** Time point 48 HPI = 242 cells.

**Figure S3. Percent of intracellular *Toxoplasma gondii* infection per cell during time-course infection in experiment #2.** There are six time points: 1, 6, 9, 12, 24, and 48 HPI. Each time point is represented by one graph. X-axis represents the percentage of intracellular parasites. The Y-axis represents the cell count, number of cells in each specific percentage. **(A)** Time point 1 HPI = 131 cells. **(B)** Time point 6 HPI = 75 cells. **(C)** Time point 9 HPI = 98 cells. **(D)** Time point 12 HPI = 81 cells. **(E)** Time point 24 HPI = 76 cells. **(F)** Time point 48 HPI = 70 cells.

**Figure S4. Establishing a threshold of *T. gondii* infection. (A)** Representative images of infected HFF with lower than 5% (yellow) and higher (red) than 5% mCherry *T. gondii*. Scale bar = 50 µm. **(B)** ORR = intensity of NAD(P)H / (intensity of NAD(P)H + intensity of FAD)**. (C)** Optical Redox ratio of *T. gondii* infected HFF cells with low vs. high intracellular *Toxoplasma gondii* in time course infection. **(D)** Percentage of protein bound NAD(P)H of *T. gondii* infected HFF cells with low vs. high intracellular *Toxoplasma gondii* in time course infection. **(E)** Mean lifetime of NAD(P)H of *T. gondii* infected HFF cells with low vs. high *Toxoplasma gondii* in time course infection. The percentage of intracellular mCherry *T. gondii* lower than 5% is represented in yellow and higher than 5% is represented in red. Values represent the median with 1.5 IQR. Statistical significance was determined by an independent Student’s T-test. *p < 0.05.**p < 0.01, ***p < 0.001, ****p < 0.0001. Cell count low *T. gondii* = 632, cell count high *T. gondii* = 974

**Figure S5. Temporal changes in optical redox ratio, NAD(P)H lifetime, and FAD lifetime of *T. gondii* infected HFF cells**. **(A)** Optical redox ratio of *T. gondii* infected HFF cells vs. uninfected cells in time course. **(B)** NAD(P)H mean lifetime (τ_m_) of *T. gondii* infected HFF cells. **(C)** Percentage of protein bound NAD(P)H (α_2_) of *T. gondii* infected HFF cells. **(D)** Percentage of protein bound FAD (α_1_) of *T. gondii* infected HFF cells. Uninfected HFF cells are represented in blue and infected HFF cells are represented in red. N = 243, 225, 285, 277, 263, 312 for 1, 6, 9, 12, 24 and 48 HPI respectively. Each point represents the average of two independent experiments. Statistical significance was determined by an independent Student’s T-test. *p < 0.05.**p < 0.01, ***p < 0.001, ****p < 0.0001. ps, picoseconds.

**Figure S6. Temporal changes in FAD lifetime of *T. gondii* infected HFF cells over 48-hrs. (A)** Representative images of FAD mean lifetimes (τ_m_) reported in ps, picoseconds. Scale bar = 50 µm. **(B-C).** FAD τ _m_ of *T. gondii* infected HFF cells as bar graphs and line plots respectively. Error bars in bar graph represent 1.5 IQR and 95% confidence interval in the line plots. Uninfected HFF cells are represented in blue and infected HFF cells are represented in red. n = 243, 225, 285, 277, 263, 312 for 1, 6, 9, 12, 24 and 48 HPI respectively. Each point represents the average of two independent experiments. Statistical significance was determined by an independent Student’s T-test. *p < 0.05.**p < 0.01, ***p < 0.001, ****p < 0.0001. ps, picoseconds

**Figure S7. Temporal changes in OMI parameters during *T. gondii* kiss and spit. (A)** Comparison of temporal optical redox ratio changes across all conditions normalized to the control of uninfected cells. **(B-D)** Show trends during *T. gondii* kiss and spit vs. its control (cells + CD). **(B)** Temporal optical redox ratio changes. **(C)** Temporal changes in percentage of protein bound NAD(P)H (α_2_). **(D)** Temporal changes in mean lifetime of NAD(P)H (τ_m_). Uninfected cells are represented in blue bars, infected cells are represented in red bars, cells treated with the inhibitor Cytochalasin D (CD) are represented in purple, and kiss and spit are represented in cyan. Each bar represents the average of two independent experiments. Statistical significance was determined by an independent Student’s T-test. *p < 0.05.**p < 0.01, ***p < 0.001, ****p < 0.0001. ps, picoseconds.

**Figure S8. Temporal changes in OMI parameters during control, *T. gondii* infection, and kiss and spit infection. (A-B)** Comparison of temporal optical redox ratio changes normalized to uninfected control cells shown in box plot and line plot respectively. **(C-D)** Temporal changes in percentage of protein bound NAD(P)H (α_2_) shown in a box plot and line plot respectively. **(E-F)** Temporal changes in NAD(P)H mean lifetime (τ_m_) shown in a box plot and line plot respectively. Uninfected cells are represented in blue, infected cells are represented in red, cells treated with the inhibitor Cytochalasin D (CD) are represented in purple, and kiss and spit are represented in cyan. Each bar represents the average of two independent experiments. Statistical significance was determined by an independent Student’s T-test. *p < 0.05.**p < 0.01, ***p < 0.001, ****p < 0.0001. ps, picoseconds.

**Figure S9. Gene expression of both host and parasite enzymes involved in glycolysis in *T. gondii* over 48-hours of infection**. The line graphs represent mRNA abundance for the host (grey) and *T. gondii* (red). Host expression is shown as fold change (infected/uninfected) on the left Y-axis and *T. gondii* expression is shown in Fragments Per Kilobase of transcript per Million mapped reads (FPKM) values in the right Y-axis, X-axis shows hours post infection. Data extracted from our previous publication [5].

**Figure S10. Host go term molecular function clustering of significant upregulated genes during ME49 *T. gondii* infection.** Data extracted from our previous publication [5]. Each of the seven time points are represented by a different color. Analysis was performed in the base data DAVID.

**Figure S11.** Host reactome clustering of significant upregulated genes during ME49 ***T. gondii* infection.** Data extracted from our previous publication [5]. Each of the seven time points are represented by a different color. Analysis was performed in the base data DAVID.

**Figure S12.** Heatmap of host enzymes gene expression most likely to bind to NAD(P)H during ME49 *T. gondii* time course infection. Data extracted from our previous publication [5]. Color scale represents the fold change with respect to the uninfected control.

**Figure S13. Expression of 58 genes related to redox biology in *T. gondii.*** Data extracted from our previous publication [5]. The color scale represents the abundance in fragments per kilobase of exon per million mapped fragments (FPKM).

**Figure S14. Expression of other genes related to redox biology in *T. gondii.*** Data extracted from our previous publication [5]. The color scale represents the abundance in fragments per kilobase of exon per million mapped fragments (FPKM).

## ACKNOWLEDGEMENTS

The authors would like to thank Cerise Siamof, Kelsey Tweed and Andres Tibabuzo for their assistance with the processing of cell masks; Amani Guillette for assistance in seahorse analysis; Dr. John Boothroyd for providing the parental RHΔKU80 and the modified RHΔKU80ΔMAF strains; Matthew Stefely for edition of figures and visualization and Alicia Williams for editing of the manuscript.

## CODE AND DATA AVAILABILITY

Codes used in this manuscript are available in https://github.com/skalalab/gallego_g-omi_toxoplasma_redox_ratio. Images are available upon request.

## AUTHOR CONTRIBUTIONS

Conceptualization, G.M.G.L., M.S. and LJK; Data curation, E.C.G.; Formal analysis, G.M.G.L. and E.C.G; Funding acquisition, M.S. and LJK.; Investigation, G.M.G.L.; Methodology, G.M.G.L. and E.C.G.; Supervision, M.S. and LJK; Validation, G.M.G.L.; Visualization, E.C.G.; Writing—original draft, G.M.G.L. and E.C.G.; Writing—review and editing, M.S. and LJK. All authors have read and agreed to the published version of the manuscript.

