## Supplementary figures and images for "Metabolic changes to host cells with *Toxoplasma gondii* infection"

### supplemental Figure 1

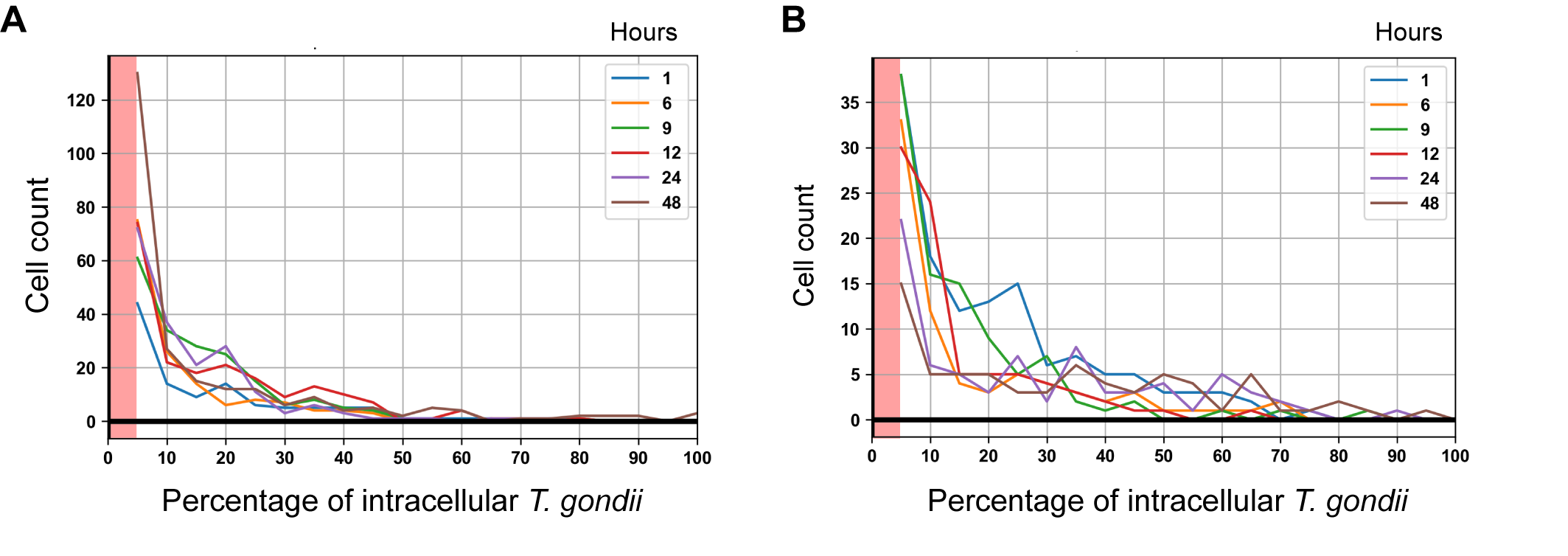

### supplemental Figure 2

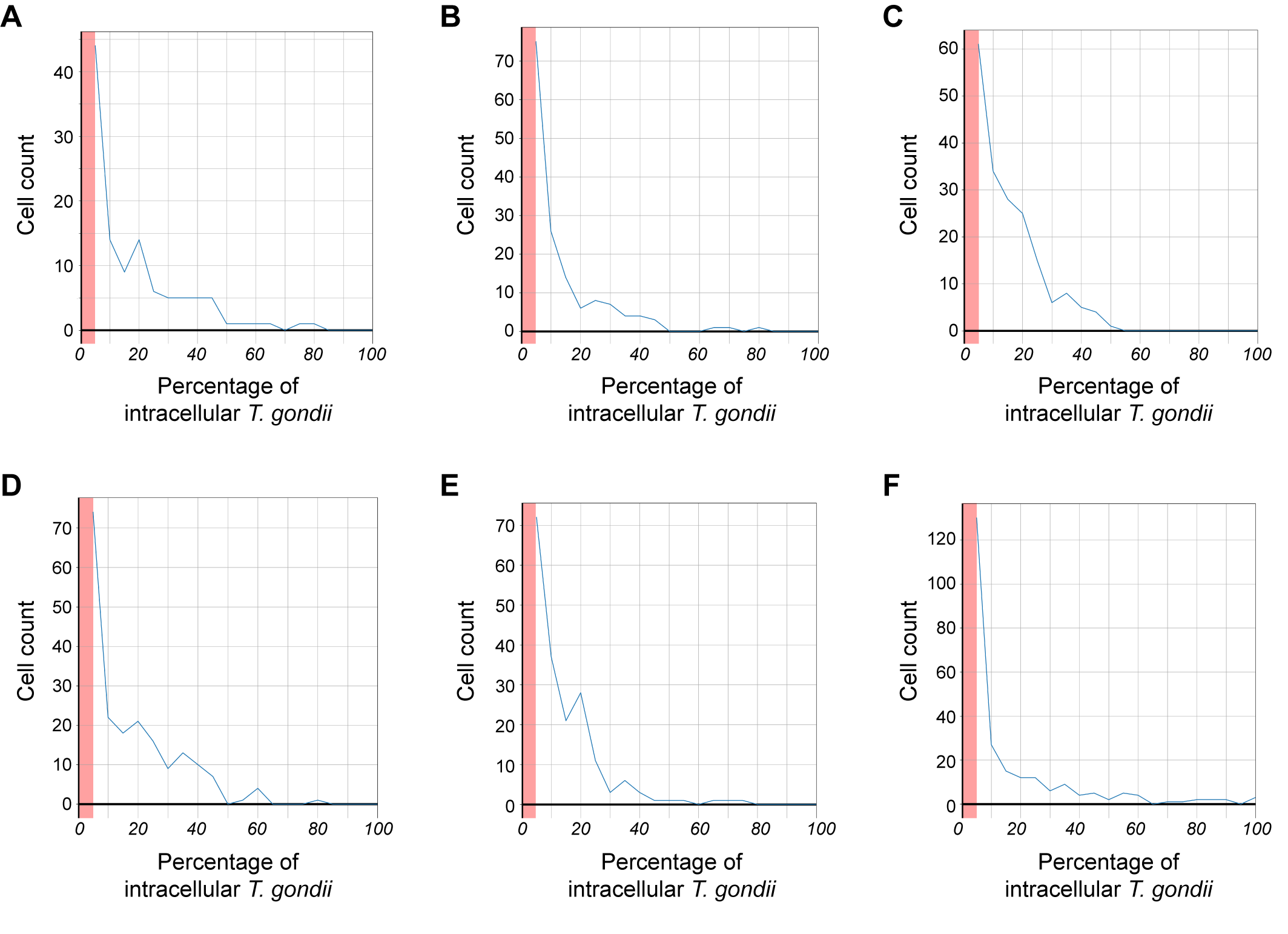

### supplemental Figure 3

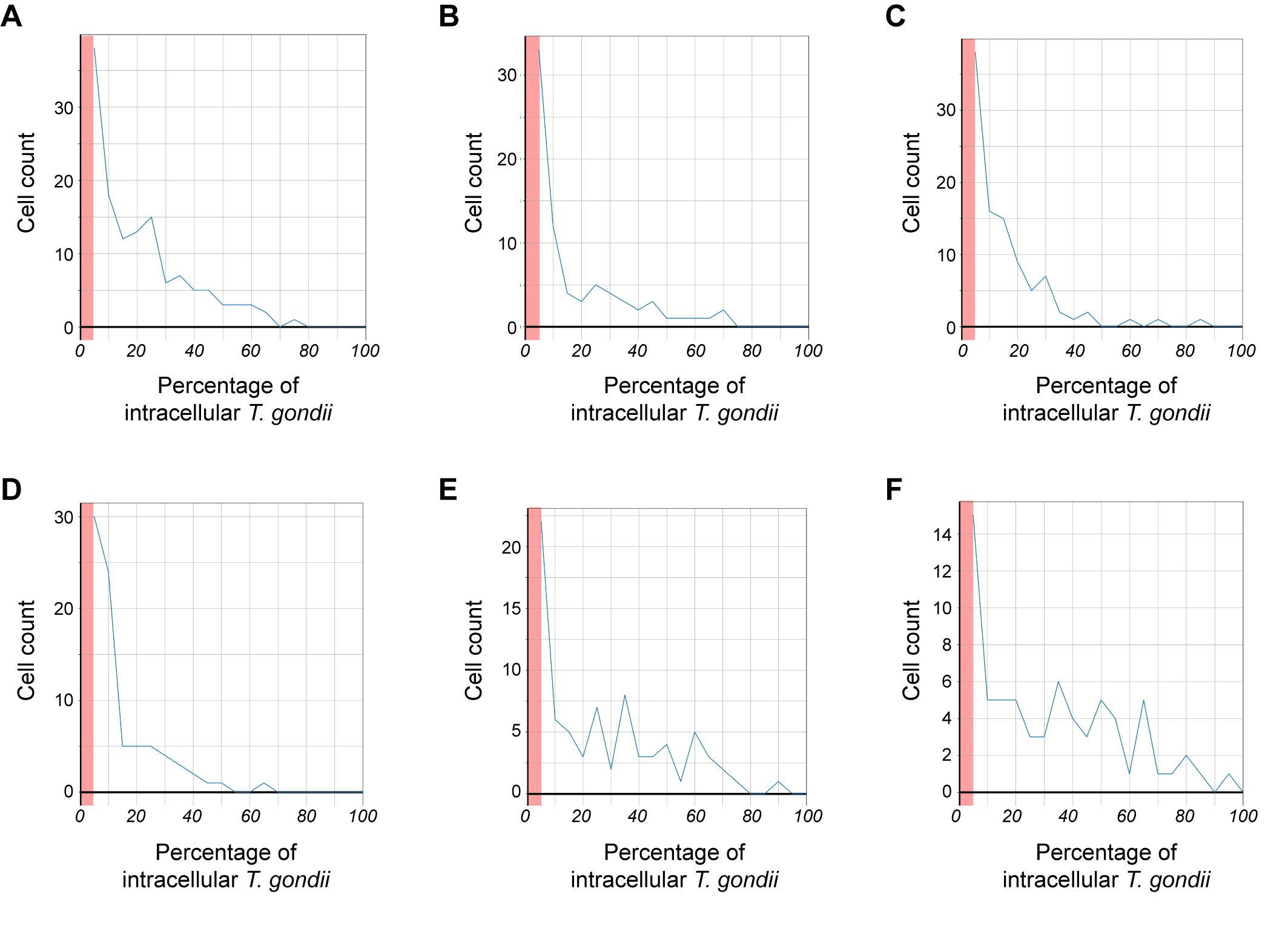

### supplemental Figure 4

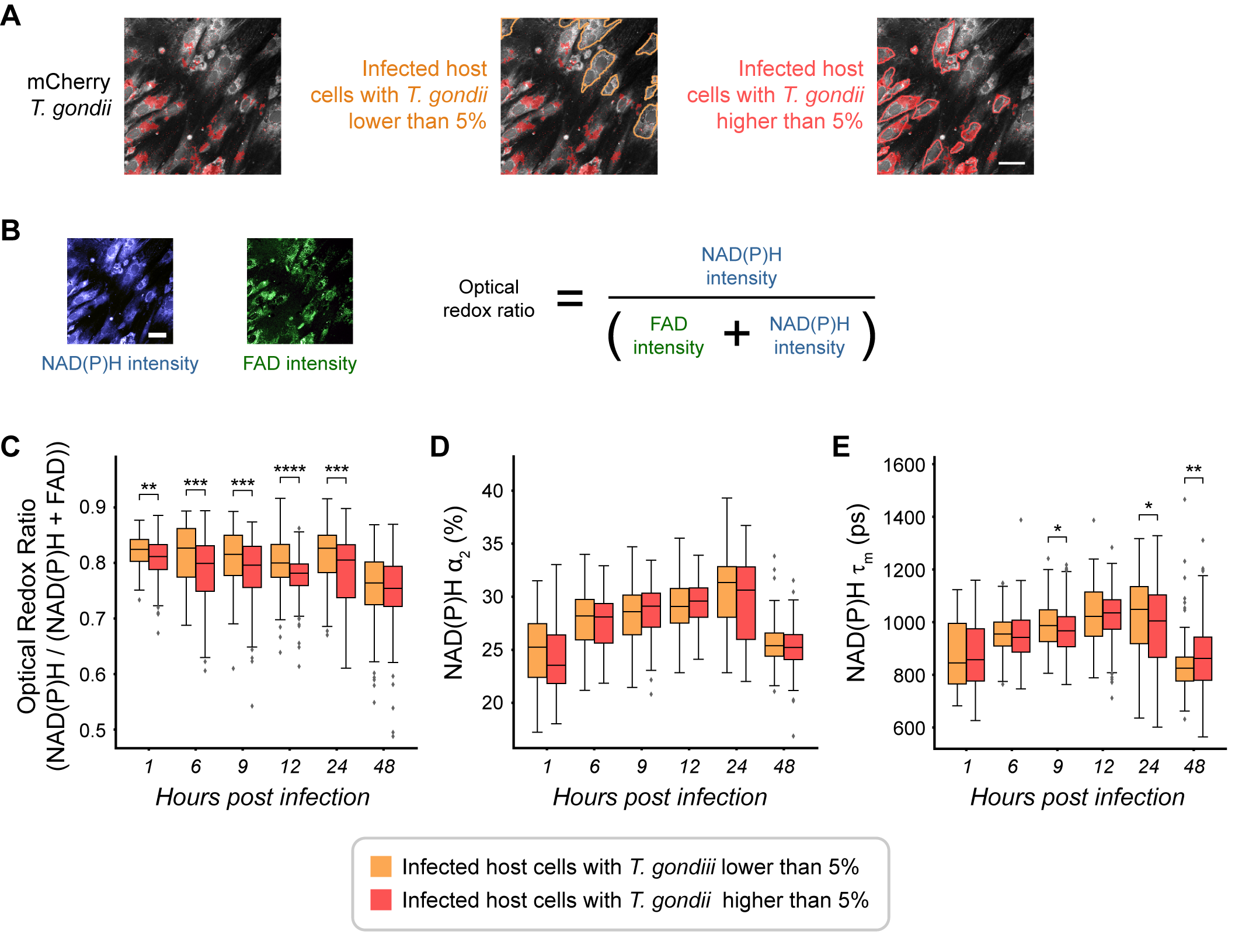

### supplemental Figure 5

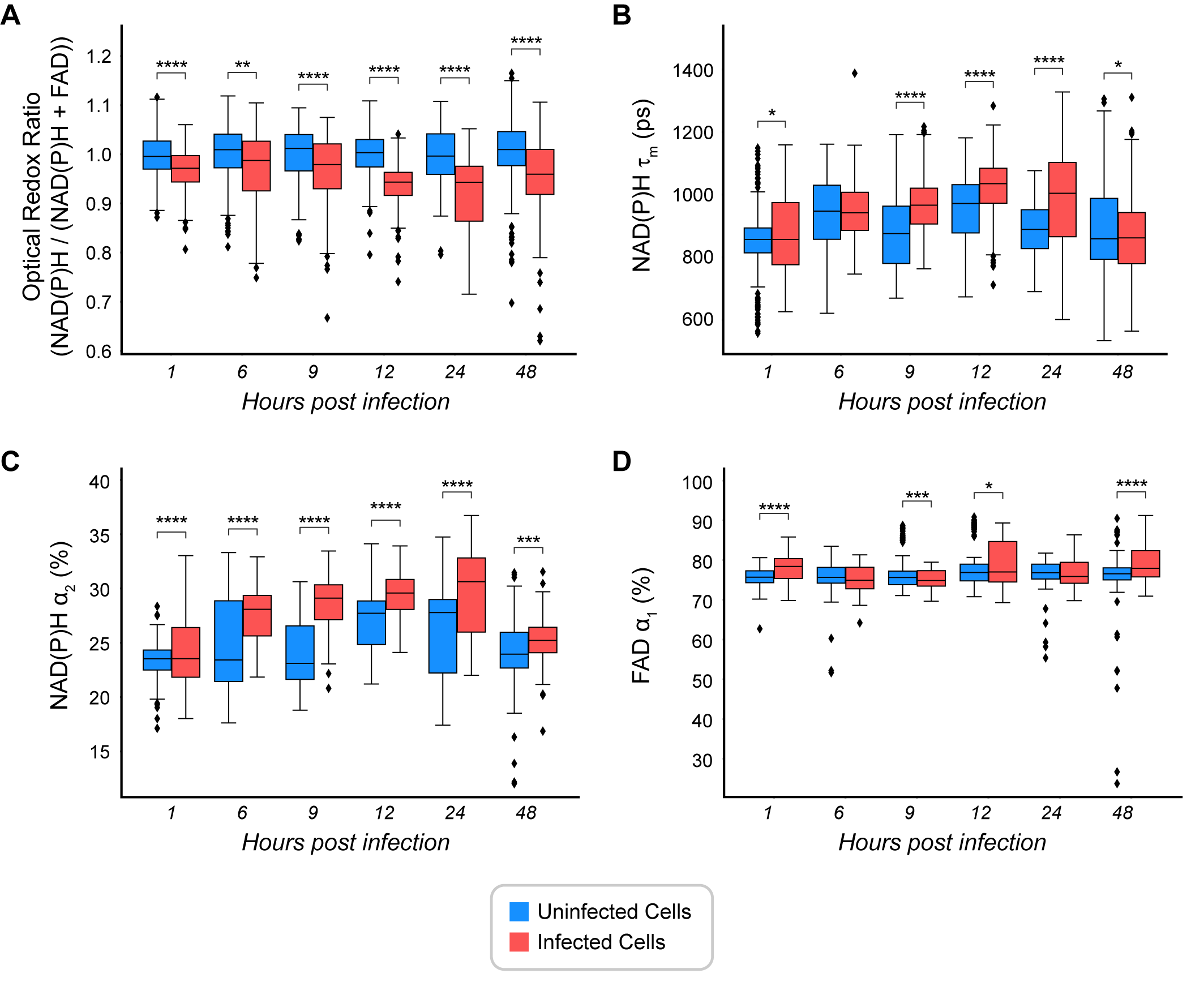

### supplemental Figure 6

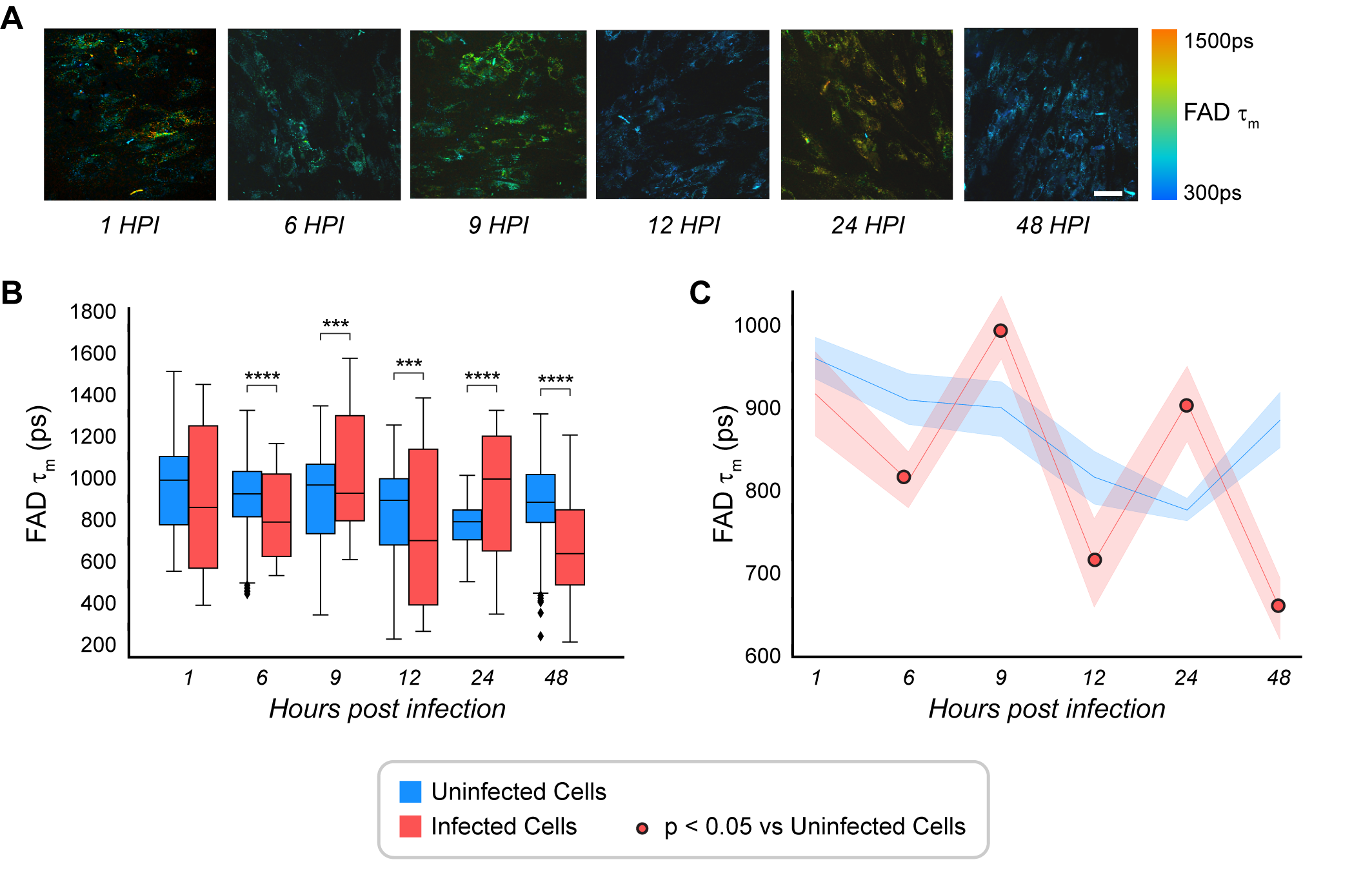

### supplemental Figure 7

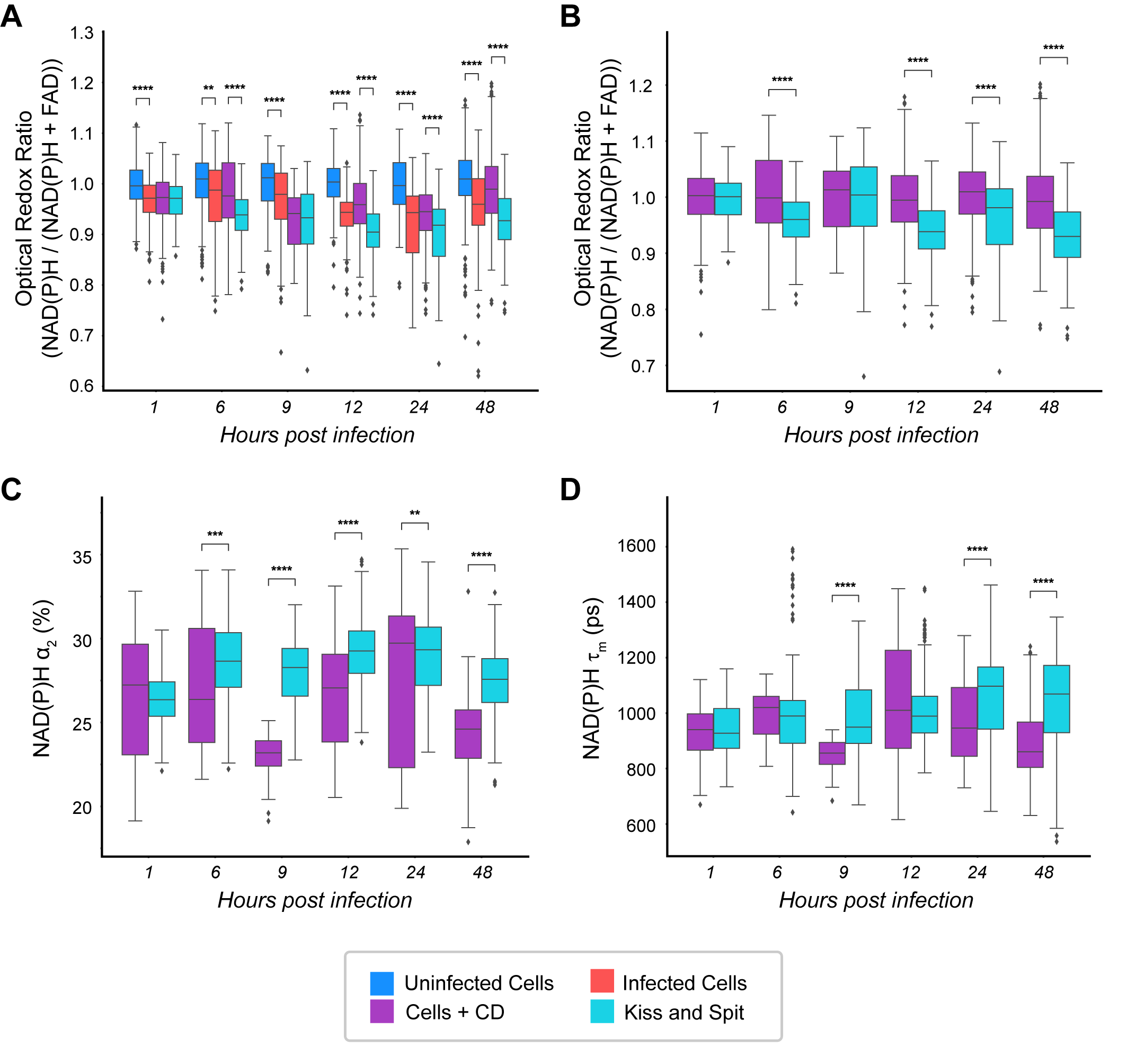

### supplemental Figure 8

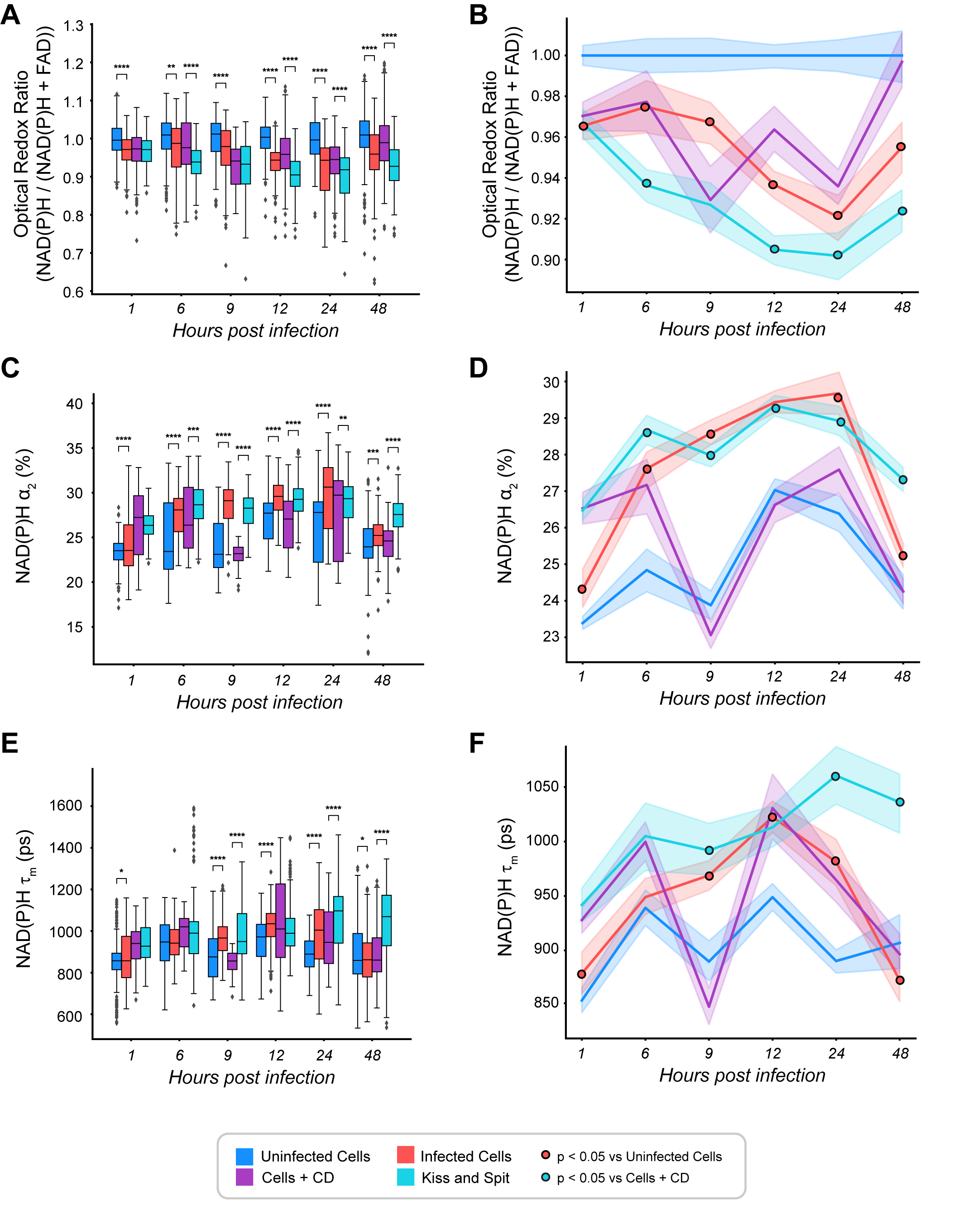

### supplemental Figure 9

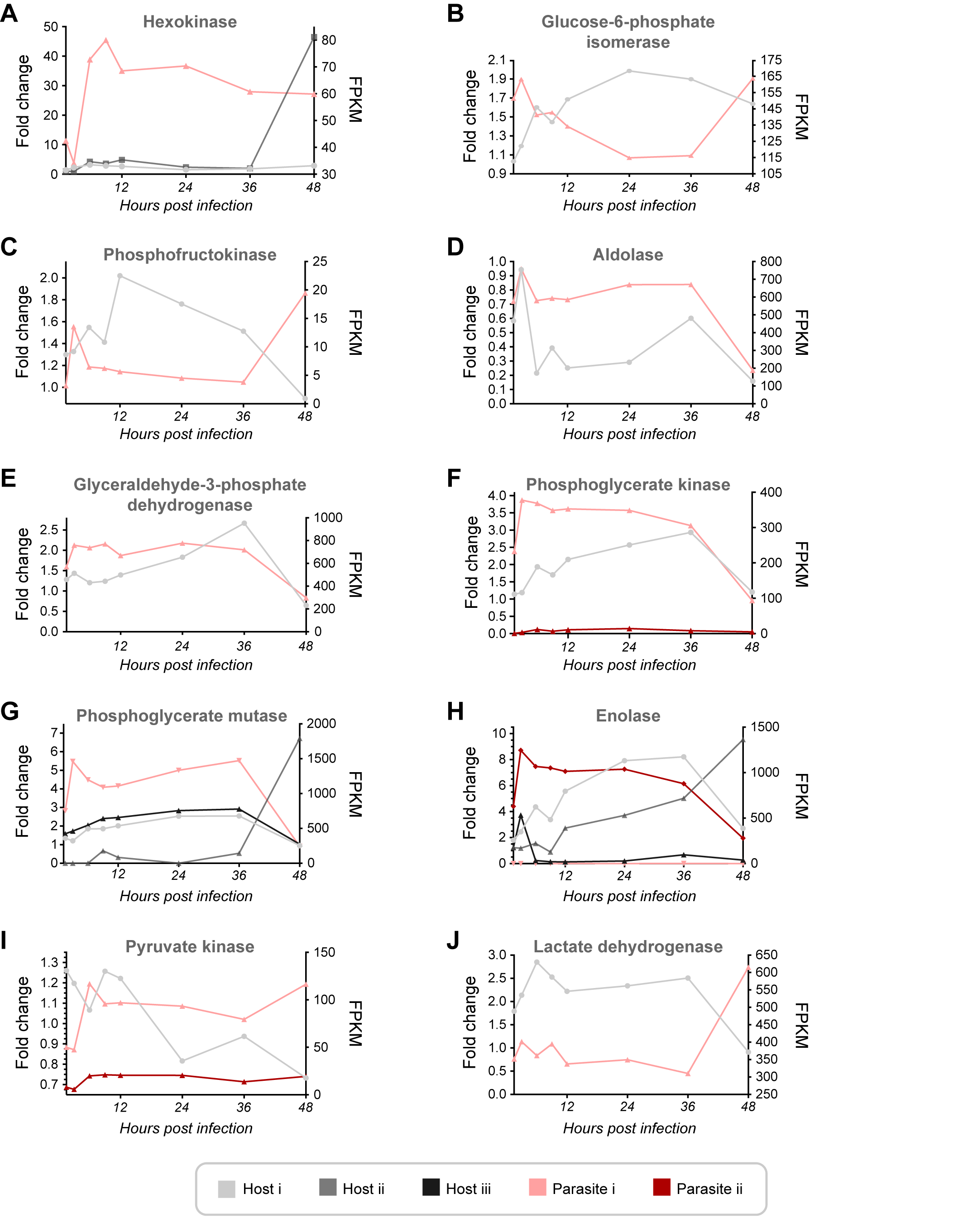

### supplemental Figure 10

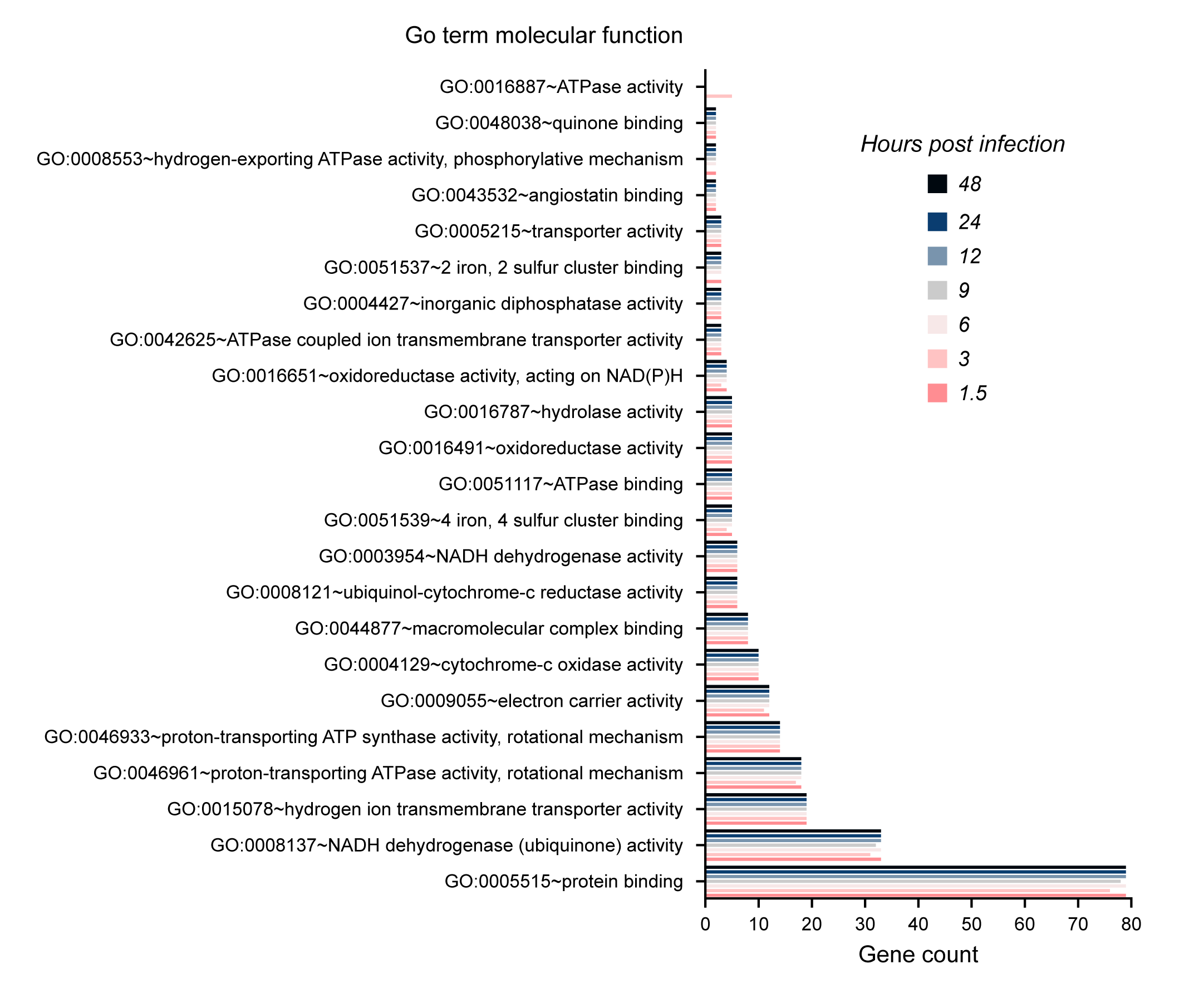

### supplemental Figure 11

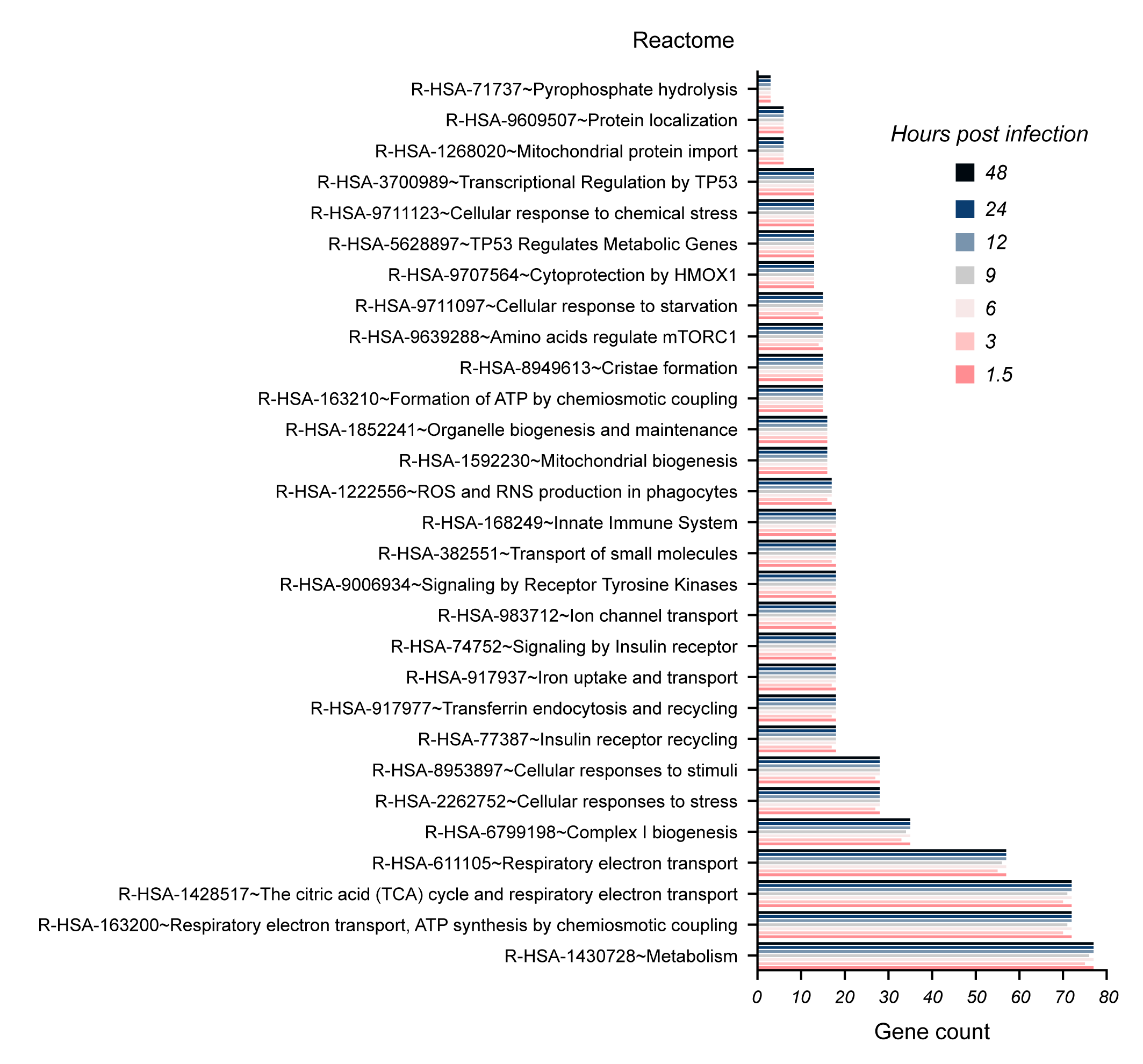

### supplemental Figure 12

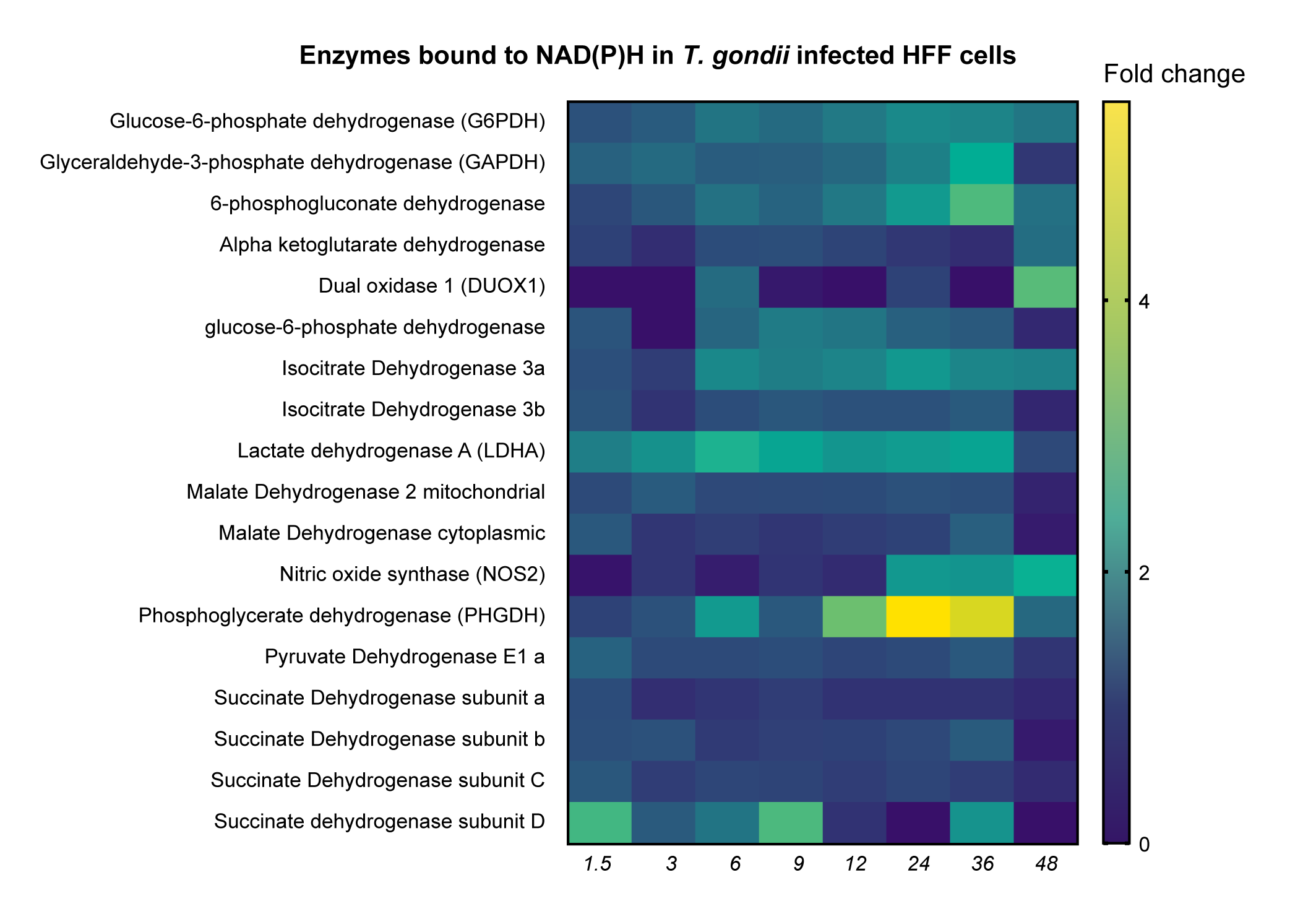

### supplemental Figure 13

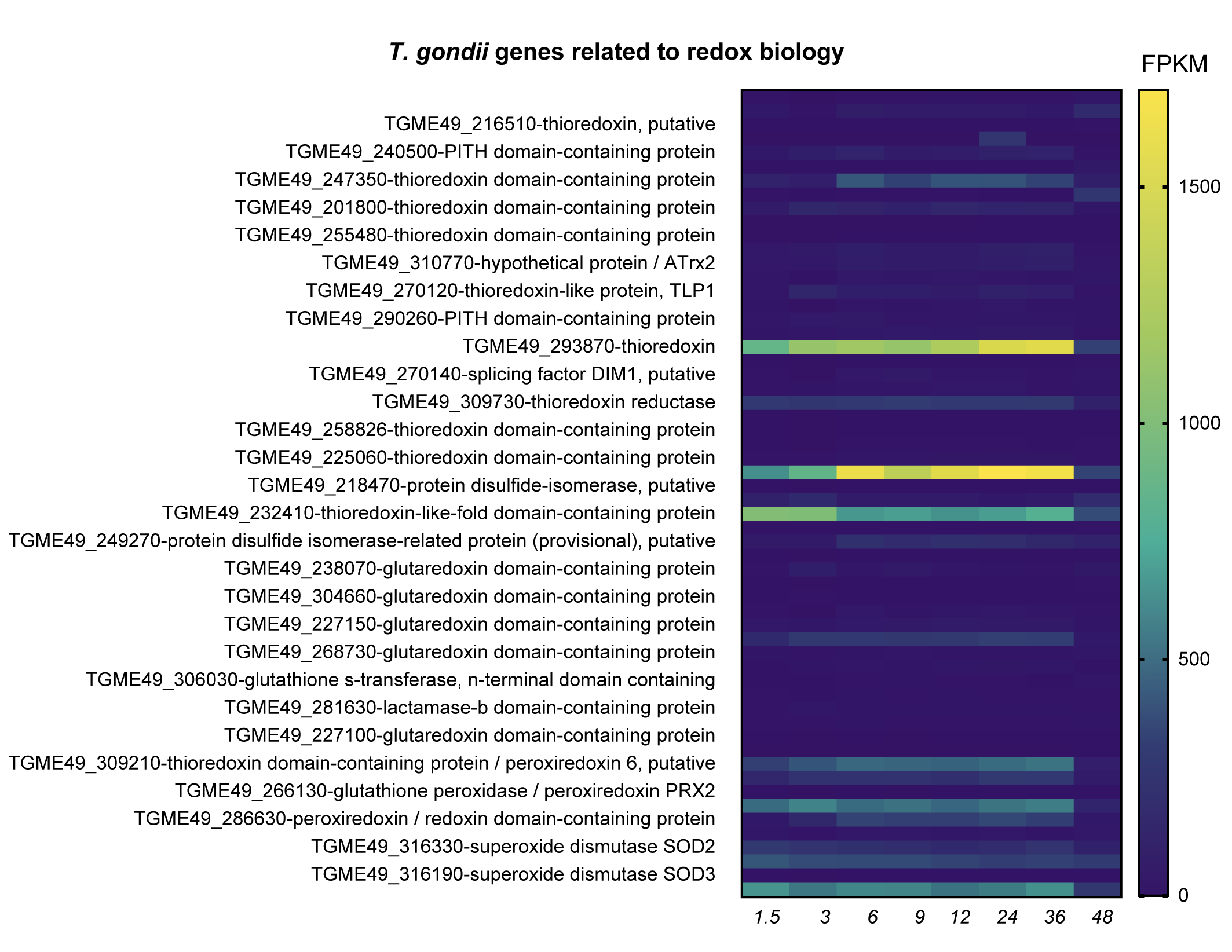

### supplemental Figure 14

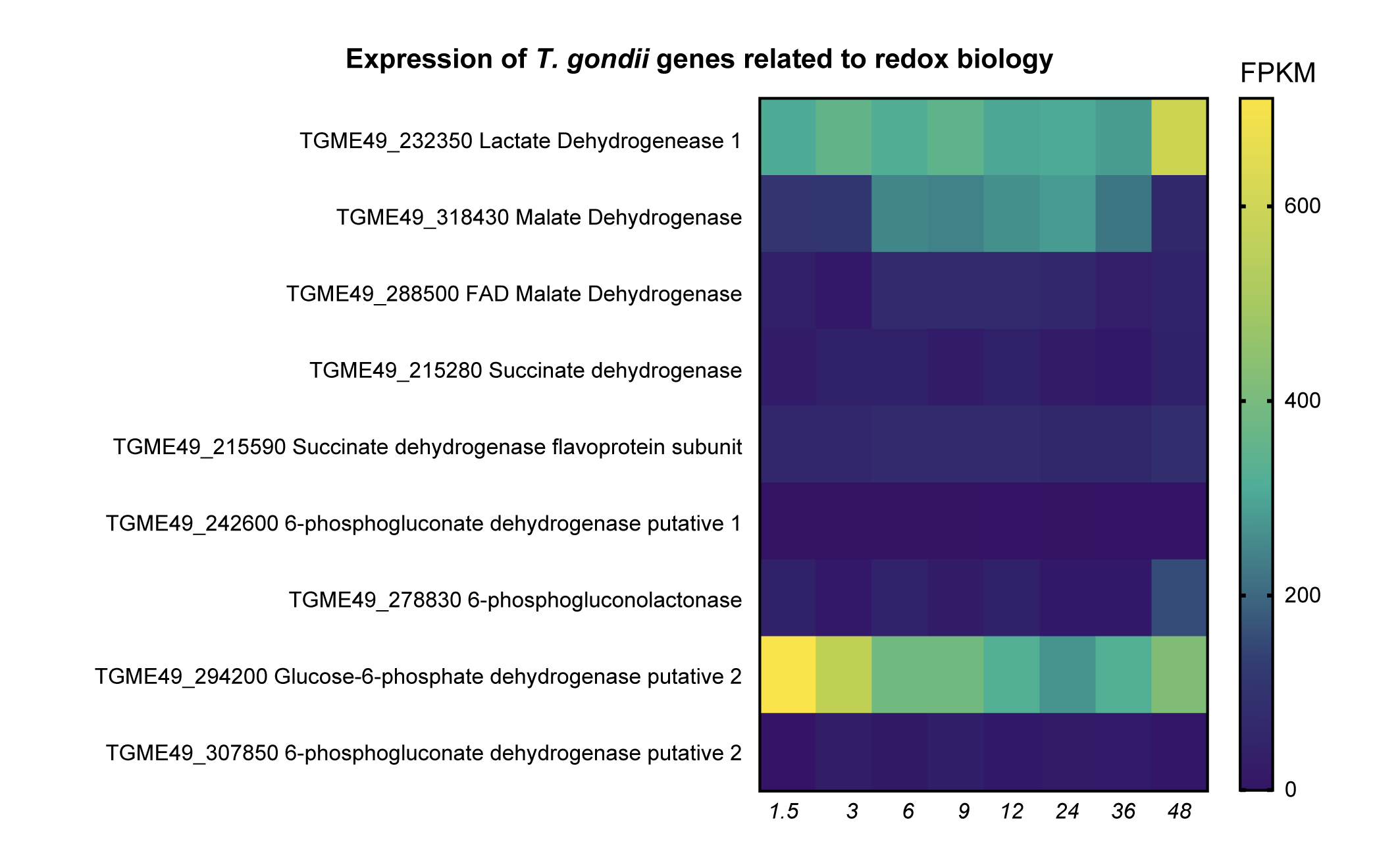
